# SCRINSHOT, a spatial method for single-cell resolution mapping of cell states in tissue sections

**DOI:** 10.1101/2020.02.07.938571

**Authors:** Alexandros Sountoulidis, Andreas Liontos, Hong Phuong Nguyen, Alexandra B. Firsova, Athanasios Fysikopoulos, Xiaoyan Qian, Werner Seeger, Erik Sundström, Mats Nilsson, Christos Samakovlis

## Abstract

Changes in cell identities and positions underlie tissue development and disease progression. Although, single-cell mRNA sequencing (scRNA-Seq) methods rapidly generate extensive lists of cell-states, spatially resolved single-cell mapping presents a challenging task. We developed SCRINSHOT (<u>S</u>ingle <u>C</u>ell <u>R</u>esolution <u>IN</u> <u>S</u>itu <u>H</u>ybridization <u>O</u>n <u>T</u>issues), a sensitive, multiplex RNA mapping approach. Direct hybridization of padlock probes on mRNA is followed by circularization with SplintR ligase and rolling circle amplification (RCA) of the hybridized padlock probes. Sequential detection of RCA-products using fluorophore-labeled oligonucleotides profiles thousands of cells in tissue sections. We evaluated SCRINSHOT specificity and sensitivity on murine and human organs. SCRINSHOT quantification of marker gene expression shows high correlation with published scRNA-Seq data over a broad range of gene expression levels. We demonstrate the utility of SCRISHOT by mapping the locations of abundant and rare cell types along the murine airways. The amenability, multiplexity and quantitative qualities of SCRINSHOT facilitate single cell mRNA profiling of cell-state alterations in tissues under a variety of native and experimental conditions.

## INTRODUCTION

Recent advances in single-cell RNA sequencing technologies (scRNA-Seq) enabled transcriptome analysis of individual cells and the identification of new cellular states in healthy and diseased conditions (1). These methods however fail to capture the spatial cellular organization in tissues due to cell dissociation. New spatial transcriptomic methods aim to circumvent the problem of lost cellular topology (2). They can be divided into two categories: First, targeted methods that directly detect specific mRNAs with single-cell resolution like ISS (3), MERFISH (4), osmFISH (5) and second global methods, which are based on barcode annotated positions and next generation sequencing to resolve RNA topology. The spatial resolution of global methods is still larger than the typical cellular dimensions (6, 7).

Targeted methodologies are based on nucleic acid probes (mainly DNA), complementary to the RNA species of interest, as in all *in situ* hybridization assays (8). Single-molecule fluorescence *in situ* hybridization (smFISH) is the most powerful among the spatial transcriptomic methods and has been used to supplement scRNA-Seq data with spatial information. It utilizes multiple fluorophore-labeled probes, which recognize the same RNA molecule along its length and visualize single RNA molecules as bright fluorescent dots (9, 10). Nonetheless, this method still retains some limitations such as low signal-to-noise ratio, reduced sensitivity on short transcripts, false positive signal due to unspecific-binding of the labeled probes and low capacity for multiplex detection of many RNA molecules (11–13). Multiplex detection with smFISH was initially addressed by the sequential fluorescence *in situ* hybridization (seqFISH) (14, 15) and the multiplexed error-robust FISH (MERFISH) (4). These approaches utilize sequential rounds of hybridization of FISH probes or barcode-based primary probes to detect multiple RNA species. The outstanding throughput of these methods makes them strong candidates for generation of spatial transcriptome maps in tissues. However, since the principle of these techniques is similar to smFISH, they require large number of gene-specific probes, confocal or super-resolution microscopy to deconvolve the signals and complicated algorithms for both probe design and analysis. Nevertheless, the low signal-to-noise ratio still remains a major technical challenge of these methods, especially for tissue sections with strong auto-fluorescence from structural extracellular matrix components like collagen and elastin (16). New strategies for signal amplifications, such as branched-DNA amplification (RNAScope) (17) and hybridization chain reaction (18, 19), have been recently combined with sophisticated probe design (AmpFISH) (20) to increase sensitivity and specificity of smFISH.

Padlock probes have been successfully used to detect RNA species (21). They are linear DNA molecules, with complementary arms to the target mRNA sequence and a common “backbone”. Upon hybridization with the target sequence, they can be ligated, creating circular single-stranded DNA molecules, which are used as templates for signal amplification using Φ29 polymerase-mediated rolling circle amplification (RCA) (22). RCA products are large single stranded DNA molecules containing hundreds of copies (23) of the complementary padlock-probe sequence. They can be detected with fluorophore labelled oligos, which recognize either their RNA-specific sequence or their backbone. Because each RCA-product contains hundreds of repeats of the same detected sequence, the signal-to-noise ratio increases significantly, facilitating signal detection by conventional epifluorescence microscopy. Also, multiplexity has been integrated into the method by sequencing by ligation (24, 25). Since commercial ligases such as T4 DNA ligase, T4 RNA ligase 2 and Ampligase show low activity on DNA/RNA hybrids, RNA has to be reverse-transcribed to cDNA fragments before introducing padlock probes (3). However, cDNA synthesis on fixed tissue sections is a challenging and expensive procedure (26), prompting new elegant approaches trying to circumvent reverse-transcription by introducing Click-chemistry to ligate DNA probes after hybridization on their RNA targets (ClampFISH (27)). More recently, “H-type” DNA probes, which are hybridized to both RNA and padlock probes (PLISH (28)), or SNAIL-design probes have been successfully used to facilitate intramolecular ligation of the padlock probes (STARmap (25)).

PBVC-1 DNA ligase (also known as SplintR ligase) shows strong ligase activity of DNA sequences in DNA/RNA hybrids. This enzyme is a DNA ligase encoded by *Paramecium bursaria Chlorella* virus 1, first discovered by Ho et al.,(29). A number of studies have successfully applied SplintR ligase for RNA species detection in cultured cells (26, 30–32). However, the fidelity of SplintR ligation in end-joining is questionable because it tolerates mismatches at the padlock probe junction and its usefulness is debated. In addition, SplintR ligase shows 1% of its ligation activity on a 1-nucleotide gap of a nicked duplex DNA substrate (26, 33).

We present SCRINSHOT an optimized protocol for multiplex RNA *in situ* detection on paraformaldehyde-fixed (PFA) tissue sections. We validated the sensitivity, specificity and multiplexity of SCRINSHOT and showed that it is quantitative over a broad range of gene expression levels. It is based on the *in situ* sequencing protocol (3) but bypasses the costly and inefficient reverse transcription on fixed tissue to gain higher detection efficiency, by utilizing SplintR ligase. To minimize false positive artifacts, we utilized 40nt long target-specific sequences in the padlock probes in combination with stringent hybridization conditions. SCRINSHOT performs on a variety of tissues including lung, kidney and heart and readily detects several epithelial, endothelial and mesenchymal cells. We tested the multiplexing of SCRINSHOT on mouse and human tissue sections, by simultaneous detection of characterized cell-type-selective markers. SCRINSHOT successfully identified distinct cell-types and helped to create spatial maps of large tissue areas at single cell resolution.

## RESULTS AND DISCUSSION

### SCRINSHOT overview

SCRINSHOT evolved from our attempts to improve the detection sensitivity and reduce the cost of the *in situ* sequencing method, using PFA-fixed material. PFA fixation significantly improves the histology but makes RNA less accessible to enzymes and padlock probes (24, 26). We focused on stringent padlock probe hybridization to RNA targets and omitted the inefficient *in situ* cDNA synthesis step (Figure 1 and Supplementary Figure 1). Gene-specific padlock probes are first directly hybridized to 40nt long, unique sequences on the mRNA targets. Upon ligation of bound padlock probes, we amplify the padlock probe sequence using Φ29 polymerase dependent rolling cycle amplification. RCA products are subsequently detected by fluorophore-labelled oligos which recognize the gene specific part of the padlock, as previously described (3). Multiplexity is reduced compared to sequencing by ligation (24), which theoretically allows the detection of 256 transcript species in 4 hybridization runs, or to other barcode based approaches (4, 26). To increase the number of detected genes we used sequential hybridization cycles of fluorophore-labelled, uracil-containing oligos. After each detection cycle, fluorescent probes are removed by enzymatic fragmentation by uracil-N-glycosylase (UNG) and stringent washes. Sequential detection also enables the separate detection of low, medium and high abundant RNA species, increasing the dynamic range of detection (34). Our detection probes were each labelled by one of three commonly-used fluorophores and allowed up to 10 hybridization and imaging cycles, typically detecting 30 genes. After image acquisition, we utilized manual segmentation of nuclei and open access analysis tools for image stitching and signal quantification to construct a simple pipeline for quantitative mapping of expression counts for 20-30 genes to thousands of cells on tissue sections. The detailed protocol starting with tissue fixation and leading to mapping is presented in the “Additional File 1”.

**Figure 1.**
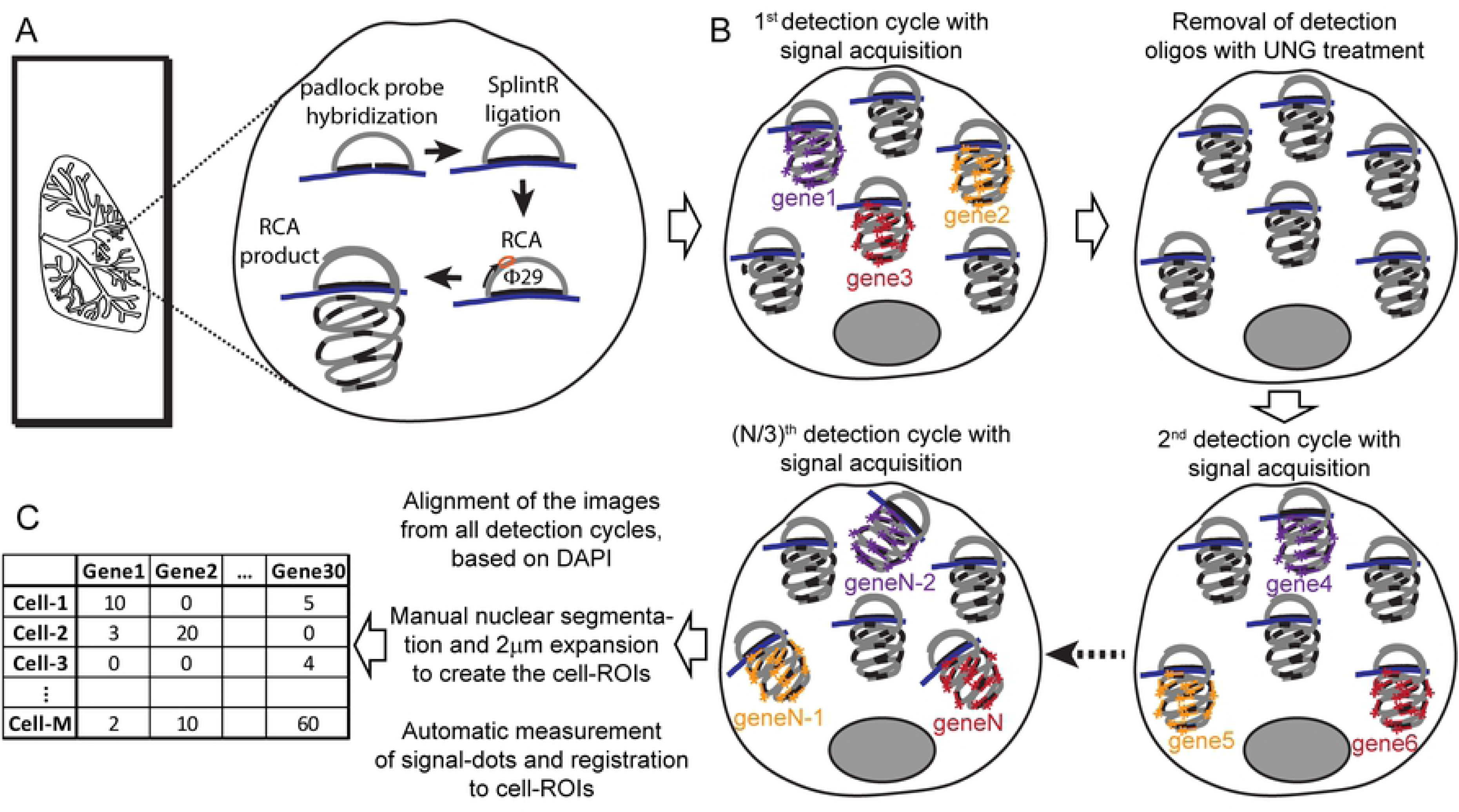
Schematic representation of SCRINSHOT. The major steps of the assay are (A) All padlock-probe hybridization for all the targeted RNA-species followed by ligation and RCA-amplification. (B) RCA-products are detected sequentially reading three per cycle, with FITC-, Cy3- and Cy5-labelled detection-oligos, which recognize the gene-specific part of the padlock probes. (C) Images from all detection cycles are aligned using DAPI nuclear staining and segmented to create the cell-ROIs and signal-dots are counted and registered to the cell-ROIs.

### SCRINSHOT specificity depends on stringent hybridization of the padlock probe

The specificity of SCRINSHOT crucially relies on the specific targeting of the SplintR ligation activity to the correct sites of the interrogated mRNA. A recent study (33) reported that the fidelity of this ligase is poor as it tolerates mismatches at padlock probe junctions. In addition, SplintR shows 1% of its ligation activity on a 1-nucleotide gap of a nicked duplex DNA substrate and it is unable to join ends across a 2-nucleotide gap (35). These results question the specificity of SplintR ligase-based methods for *in situ* RNA detection. We reasoned that the choice of padlocks with high melting points (Tm around 70°C) followed by stringent washes after DNA/RNA hybridization would circumvent SplintR promiscuity. We tested the dependence of SCRINSHOT signals on the hybridization and ligation steps of the padlock, by generating mutant padlocks, predicted to affect either the hybridization of the 3’padlock arm or the sequence of the ligation site. The 3’-scrambled arm of this padlock is expected to fail in hybridizing with the *Scgb1a1* mRNA, resulting in a linear, unligated padlock and therefore in a block in RCA. The single mismatch probe contains a single replaced nucleotide at the ligation site in 5’-end (C to G). This substitution was designed to address the effect of promiscuous ligation on the signal. In the same experiment, we included a slide, where we omitted the SplintR ligase from the reaction mixture to test whether padlock hybridization alone is sufficient to generate some RCA (SplintR^neg^; Figure 2 D, D’). In all tested conditions we also used the *Actb* normal padlock probe. This transcript was detected at similar levels in all slides and served as a control for the reactions with the mutated padlocks. We first counted the dots of *Scgb1a1* and *Actb* signals in all airway cells of sequential lung tissue sections and plotted their ratios at the different conditions (Figure 2). The signal from single-mismatch padlock probe was reduced by 20%, compared to the normal *Scgb1a1* probe demonstrating the low SplintR fidelity for the sequence of the ligation site (Figure 2 B, B’, E). The *Scgb1a1* signal was lost when we used the 3’-scrambled padlock, indicating that padlock probe hybridization is necessary for circularization and subsequent RCA (Figure 2 C, C’, E). The omission of SplintR from the ligation mixture resulted in undetectable signal for both *Scgb1a1* and *Actb* indicating the central role of ligation in signal amplification (Figure 2 D, D’, E). We noticed that the *Scgb1a1* signal showed significant crowding and a potential saturation leading to underestimation of the total number of RCA-products for this highly expressed gene (see below). This crowding was evident even when we added 5-fold less *Scgb1a1* padlock probe (0.01 μM) to the reactions in comparison to all other padlock probes. In an attempt to more accurately quantify the differences between the signals from normal and mutated padlocks we measured the overall fluorescence intensity (Raw Integrated Density) of the airway cell-ROIs. This showed that the single mismatch at the ligation site of *Scgb1a1* padlock probe causes 3-fold fluorescence signal reduction arguing that the *in situ* SplintR activity is substantially reduced, but not abolished by single nucleotide substitutions at the ligation site (Figure 2F). We conclude that the specificity of SCRINSHOT assay is largely provided by the hybridization stringency of the padlock probes, since SplintR is unable to ligate off-target padlock probes, but can ligate single-mismatch probes with low efficiency.

**Figure 2.**
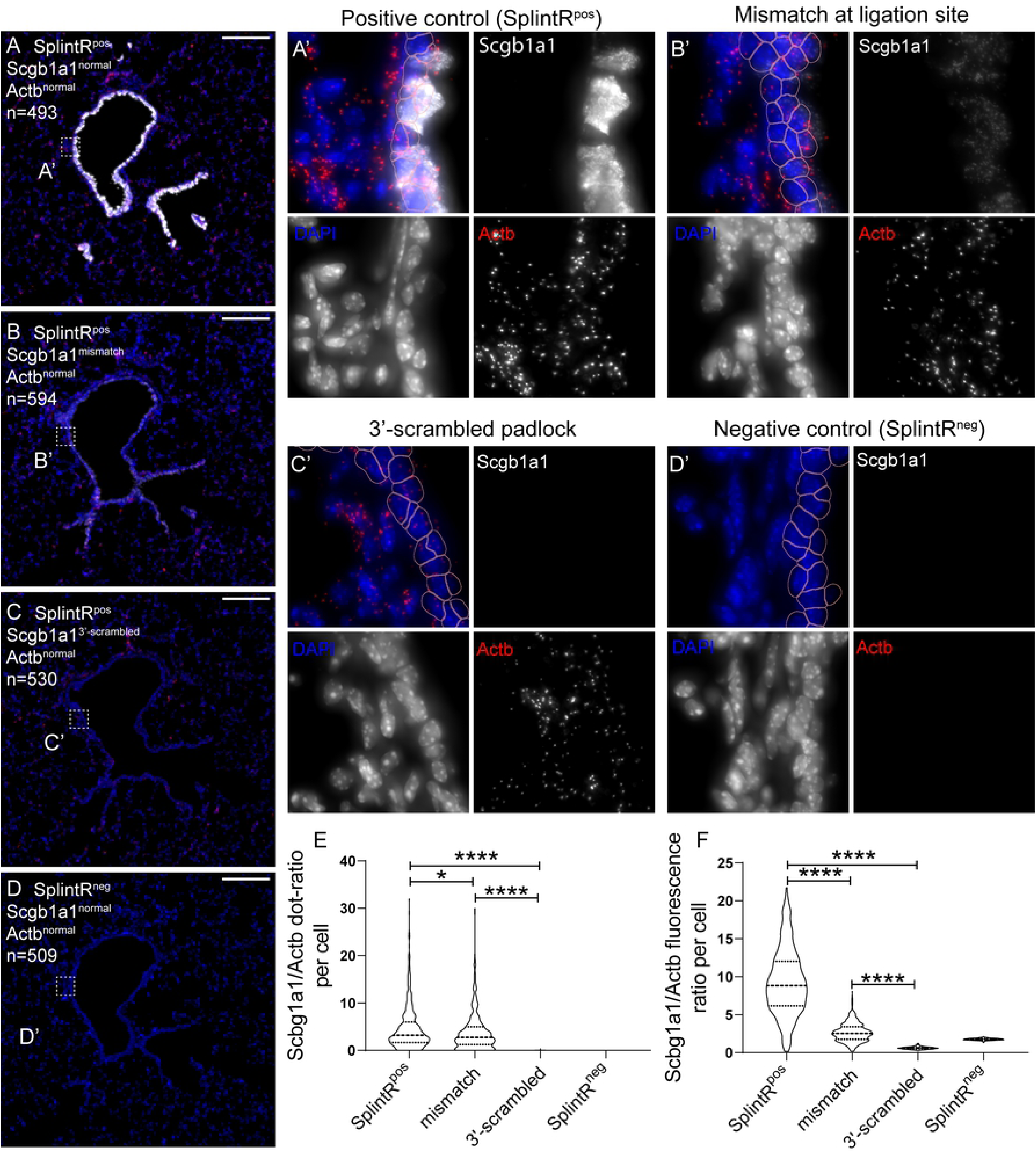
SCRINSHOT specificity relies on stringent hybridization of padlock probes to their target RNAs. Images of SCRINSHOT signal, using normal *Scgb1a1* padlock probe (A), a *Scgb1a1* padlock probe with a point mutation at its ligation site (B), a *Scgb1a1* padlock probe with 3’-scrambled arm (C) and normal padlock probe but omitting SplintR ligase (D). *Actb* normal padlock probe was used in all conditions as internal control. DAPI: blue, *Scgb1a1*: gray, *Actb*: red. “n” indicates the number of airway cells in the corresponding images. (A’-D’) Magnified areas of the indicated positions (square brackets) of images in the left. Pink outlines show the 2 μm expanded airway nuclear ROIs, which are considered as cells. Scale-bar: 150 μm. (E) Violin plot of the Scgb1a1 and Actb signal-dots ratio in all airway cells. The ratio of cells with zero *Actb*-dots considered as zero. (F) Violin plot of the *Scgb1a1* and *Actb* fluorescence intensity ratio in all airway cells. SplintR^pos^ n=473, mismatch n=574, 3’-scrambled n=507 and SplintR^neg^ n=488.

### A *Scgb1a1* antisense oligonucleotide competes with padlock probe hybridization and signal detection

If the padlock hybridization is the critical step for signal generation then competition by an oligonucleotide, which recognizes the binding to the mRNA is expected to proportionally reduce the detected RCA signal. We used the *Scgb1a1* and *Actb* padlocks together with increasing concentrations of a competing, unlabeled oligonucleotide complementary to the mRNA sequence recognized by the *Scgb1a1* padlock. Inclusion of the competitor reduced the *Scgb1a1* signal, in a dose-dependent manner. Equal molar ratios of padlock probe and competitor caused signal reduction by 10-fold and the signal was eliminated when 5-fold excess of the competitor was used (Figure 3). This suggests that the SCRINSHOT signal is proportional to the target expression levels because it can be proportionally competed with increasing concentrations of a synthetic oligo masking the hybridization site. It also highlights the importance of proper padlock design to achieve similarly high Tm values and hybridizations conditions for the different probes.

**Figure 3.**
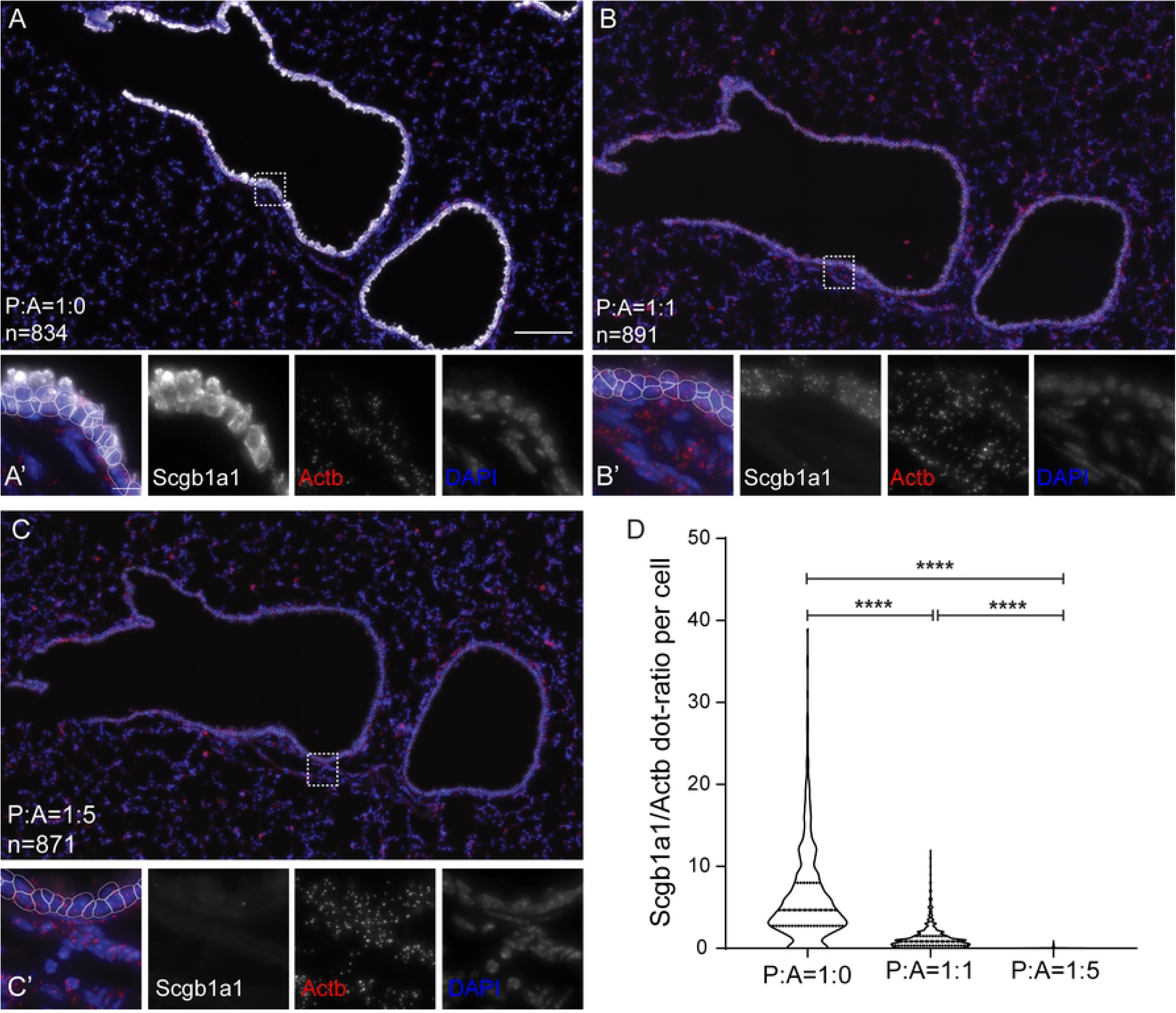
Concentration-dependent SCRINSHOT detection efficiency. (A-C) Representative images of SCRINSHOT signals for *Scgb1a1* padlock probes in the absence or presence of an antisense competitor oligo targeting the binding site of *Scgb1a1* padlock probe. SCRINSHOT signal for *Scgb1a1* padlock probe in the absence of the antisense competitor oligo (A), when mixed in equal concentration with the antisense competitor oligo (B) and when mixed with five times higher concentration of the antisense competitor oligo (C). (A’-C’) Magnified areas of the indicated positions (square brackets) of images A-C. (D) Violin plot of the *Scgb1a1* and *Actb* signal-dots ratio in all airway cells of the three compared conditions. P: padlock probe, A: antisense competitor. Scale bars in (A-C) is 100 μm and (A’-C’) is 10μm. All analyses were done using raw images and the same thresholds. For visualization purposes, brightness and contrast of *Scgb1a1* were set independently in the compared conditions to show the existence of signal in the presence antisense competitor and avoid signal saturation in its absence.

### Application of SCRINSHOT in other organs

To evaluate SCRINSHOT applicability to other tissues, we performed the assay using PFA-fixed sections from adult mouse kidney and heart and human embryonic lung. On murine tissues, we used a common panel of padlocks targeting validated lung cell type markers. We targeted *Actb* as a generic marker, *Pecam1* as an endothelial cell marker, *Scgb1a1* as a club cell marker, *Sftpc and Napsa* as alveolar epithelial type II (AT2) cell markers and *Lyz2*, as a marker for AT2 cells, macrophages and neutrophils (36). As expected, *Actb* was uniformly expressed in kidney and heart while *Scgb1a1* and *Sftpc* were undetectable (Supplementary Figure 2 A, B). In both tissues, *Lyz2* was expressed by a few scattered cells, which presumably correspond to macrophages (37, 38) (Supplementary Figure 2). In a subset of kidney tubular structures, we detected *Napsa* (Supplementary Figure 2 A), which agrees with the previously described immunohistochemical detection of the marker in renal proximal tubule cells (39). In the vessel walls of the heart, we detected sparse signal for the endothelial cell marker *Pecam1* but the myocardial cells were negative for the marker (Supplementary Figure 2 B). In the human embryonic lung sections, we used probes targeting transcripts encoding 3 transcription factors, *SOX2*, *SOX9* and *ASCL1* (Supplementary Figure 3), which have previously been detected by antibody staining in subsets of epithelial cells (40, 41). In agreement with the published results, the SCRINSHOT signal for *SOX2* was confined mainly in the proximal part of the branching epithelium, whereas *SOX9* was selectively expressed in the distal tips. *ASCL1* expression overlapped with *SOX2* (Supplementary Figure 3). These experiments show that SCRINSHOT can be readily applied to map cell-type heterogeneity in a variety of tissues.

### SCRINSHOT generates quantitative gene expression profiles in single cells

We first tested the quantitative power of SCRINSHOT by correlating its detection performance with the fluorescence of a transgenic red fluorescent protein (RFP) in mouse lung tissue sections. In the *Sftpc-CreER;Rosa-Ai14* reporter mouse, the RFP expression is activated in AT2 cells, upon Tamoxifen induction of the Cre recombinase (42). Cre recombines out a transcriptional/translational STOP cassette (43) of the Rosa26 locus and allows RFP protein expression and fluorescence in AT2 cells (44). We injected pups with Tamoxifen on postnatal day 1 (P1) and analyzed the lungs on day P21. In the same experiment, we also used lungs from *Sftpc-CreER*^neg^-*Rosa-Ai14*^pos^ mice sacrificed at P21 and lungs from *wild type* mice sacrificed at P60 as controls for the RFP induction and the potential effects of tissue autofluorescence in young and fully developed lungs.

We first analyzed 14167 *Sftpc-CreER*^pos^-*Rosa-Ai14*^pos^ cells and found a reliable correlation (R^2^=0.7233) between the endogenous RFP fluorescence and the SCRINSHOT-detected *RFP* mRNA molecules in each cell (Figure 4 B). By contrast, there was no correlation between *RFP* SCRINSHOT signal and the endogenous fluorescence (R^2^=0.0671) in 3355 *Sftpc-CreER*^neg^-*Rosa-Ai14*^pos^ cells. As expected, in the 1008 analyzed *wild type* cells, the correlation of RFP fluorescence and SCRINSHOT dots was very low. Upon closer inspection most of the signal in this lung was due to high auto-fluorescence from red blood cells, illustrating the specificity of SCRINSHOT (Figure 4 A, B).

**Figure 4.**
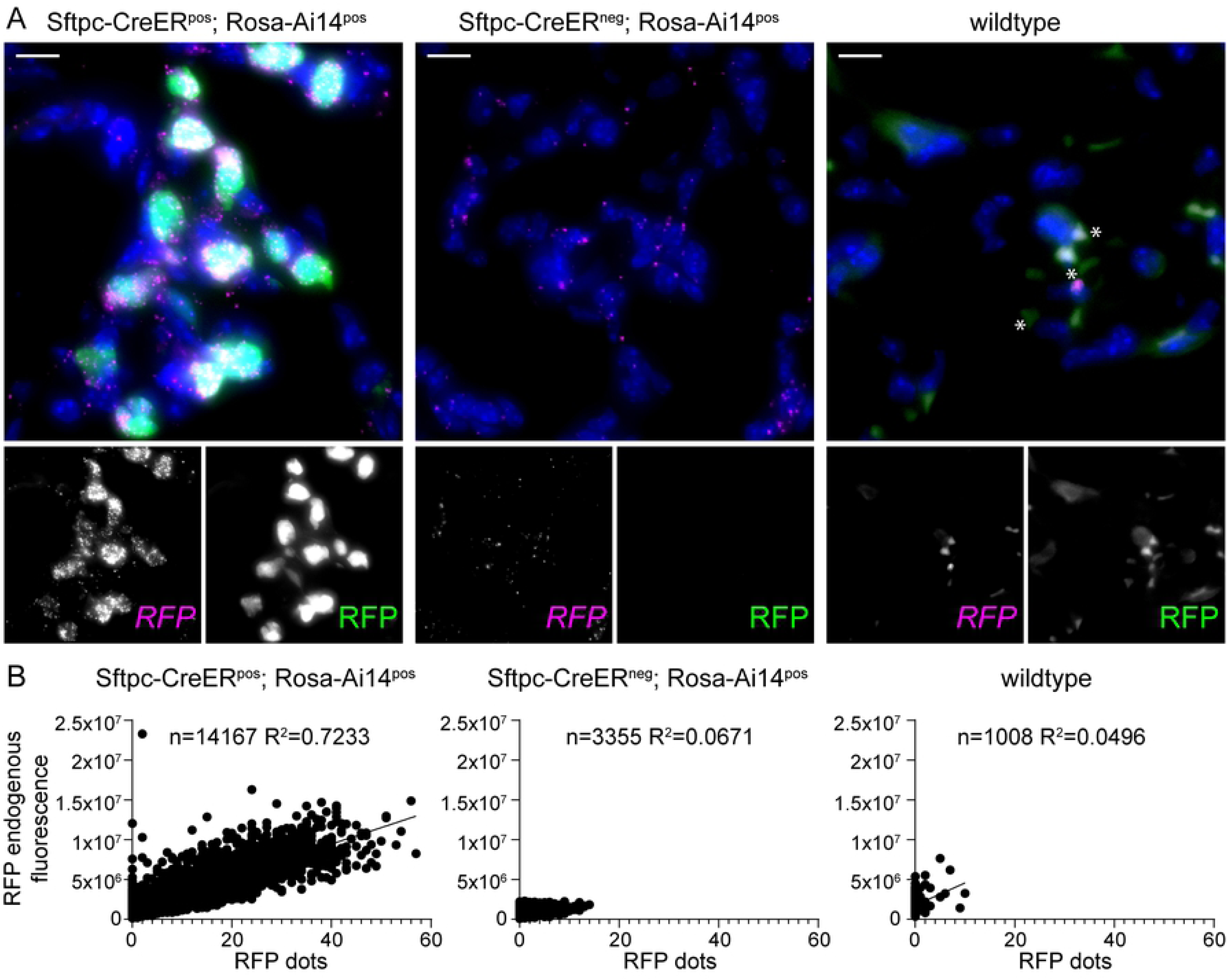
Efficient detection of *Rosa-Ai14* mRNA upon *Sftpc-CreER* driven recombination, using SCRINSHOT. (A) Representative images of alveolar regions from P21 *Sftpc-CreER*^pos^;*Rosa-Ai14*^pos^, P21 *Sftpc-CreER*^neg^;*Rosa-Ai14*^pos^ and P60 wild-type (*Sftpc-CreER*^neg^;*Rosa-Ai14*^neg^) lung sections. Asterisks indicate red blood cells with high auto-fluorescence, which give some false positive signals in wild-type lung, only. DAPI: blue, *RFP* mRNA detected with SCRINSHOT: magenta and endogenous fluorescence of RFP: green. Scale-bar: 10μm. (B) Regression analyses of RFP endogenous fluorescence (RAW integrated density) and *Ai14* SCRINSHOT dots, in the three analyzed lung sections, using Graphpad Prism. “n” indicates the number of total cells in the analyzed images, which used for the statistical analysis.

Considering that the *Sftpc-CreER;Rosa-Ai14* reporter specifically labels the alveolar AT2 cells (42), we used SCRINSHOT to identify the *Sftpc*^pos^ AT2 cells and analyze their RFP expression in both RNA (SCRINSHOT) and protein fluorescence (Raw Integrated Density) level (Figure 5 A, B, D). In *Sftpc-CreER*^pos^ cells the mean number of SCRINSHOT detected *RFP* mRNA molecules was 6-fold higher compared to *Sftpc-CreER*^neg^ cells. In the same comparison the RFP endogenous fluorescence was 5-fold higher. In *wild type* cells, the SCRINSHOT *RFP* signal was hardly detectable. We observed a slight increase in RFP fluorescence, in *wild type* cells compared to the ones from the *Sftpc-CreER^neg^* mouse, presumably due to different levels of tissue autofluorescence depending on developmental stage and fixation time of the tissues (4 hours for P21 *Sftpc-CreER* and 8 hours for *wild type* P60).

**Figure 5.**
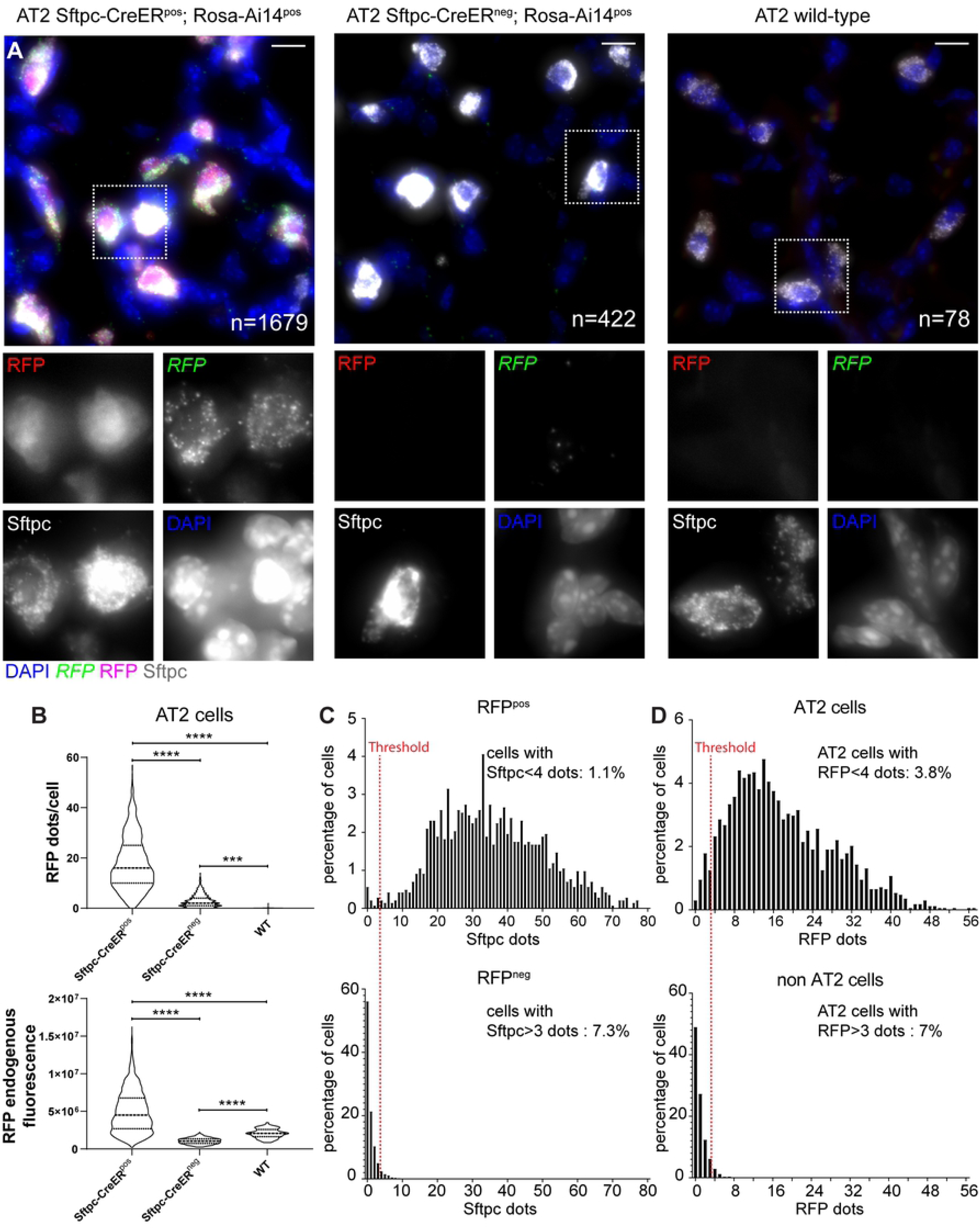
Correlation of *Sftpc* SCRINSHOT dots with RFP endogenous fluorescence and *RFP* mRNA signals. (A) Images of alveolar compartment from P21 *Sftpc-CreER*^pos^;*Rosa-Ai14*^pos^, P21 *Sftpc-CreER*^neg^;*Rosa-Ai14*^pos^ and P60 wild-type (*Sftpc-CreER*^neg^;*Rosa-Ai14*^neg^) lung sections, showing *Sftpc*^pos^ (gray) AT2 cells in correlation to *Ai14* SCRINSHOT dots (detected *RFP* mRNA) (green) and RFP endogenous fluorescence (red). Scale-bar: 10 µm. “n” designates the number of analyzed AT2 cells. (B) Violin plots of SCRINSHOT results for *RFP*, in addition to RFP endogenous fluorescence (RAW integrated density) in AT2 cells of the analyzed lungs. (E) Bar plots of SCRINSHOT results for *Sftpc* in RFP^pos^ and RFP^neg^ alveolar cells, as indicated by endogenous fluorescence of *Sftpc-CreER*^neg^;*Rosa-Ai14*^pos^ alveolar cells (threshold 2304340 units of raw integrated density), in addition to RFP in *Sftpc*^pos^ and *Sftpc*^neg^ AT2 cells. Threshold is set to 3 dots and shown with red dotted lines.

Apart from the AT2 cells, the lung alveolus contains additional cells types, including endothelial, inflammatory, epithelial AT1 cells and fibroblasts (45). In the *Sftpc-CreER*^pos^*-Rosa-Ai14*^pos^ lung, AT2 cells should be both *RFP*^pos^ and *Sftpc*^pos^ whereas the rest should be *RFP*^neg^. To evaluate the specificity and sensitivity of SCRINSHOT in the strict context of a transgenic, knock-in marker expression, we analyzed the *Sftpc* signal in *RFP*^pos^ and *RFP*^neg^ alveolar cells and the SCRINSHOT *RFP* signal in *Sftpc*^pos^ and *Sftpc*^neg^ alveolar cells from the *Sftpc-CreER*^pos^-*Rosa-Ai14*^pos^ mouse. Both genes are highly expressed, therefore we applied a threshold of 3 dots/cell to score a cell as positive (see also Supplement for details on setting detection thresholds). Of the 1429 analyzed *RFP*^pos^ cells only 1.1% did not express *Sftpc* and were thus scored as false negatives. Among the 6848 *RFP*^neg^ cells, 7.4% were positive for *Sftpc* and were considered as false positives. Conversely, we failed to detect *RFP* signal in 4% of the 1679 analyzed *Sftpc*^pos^ cells and counted them as false negatives. Among the *Sftpc*^neg^ cells 7% scored as positive for SCRINSHOT *RFP* signal and were designated as false positives (Figure 5 C, D). Thus, the levels of false positive and false negative cell annotations by SCRINSHOT are on average 5%. This is likely an overestimate because Tamoxifen-induced recombination is rarely 100% efficient and because leaky RFP transcripts from the *Rosa* locus may escape nonsense-mediated RNA decay (NMD) (46). In conclusion, the analysis of the inducible knock-in reporter shows that the SCRINSHOT *RFP* signal highly correlates with RFP endogenous fluorescence intensity in *Sftpc-CreER*^pos^ AT2 cells only. The low levels of false positive and false negative SCRINSHOT signals for *RFP* and *Sftpc* argue for the efficiency of SCRINSHOT in the identification and quantification of gene expression in alveolar cell-types.

### Multiplex performance and gene expression quantification using SCRINSHOT

To further explore the utility of SCRINSHOT in the spatial identification of cell types, we first tested if multiple rounds of hybridization and detection in lung tissue sections lead to loss of detection signal for the genes of interest, as seen in other transcriptomic approaches (5). We compared the detection signals of *Calca*, a known neuroendocrine gene marker during the first and the eighth cycles of hybridization and detection and found no significant loss of signal or decreased specificity (Supplementary Figure 4). In the same experiment, we also detected a pair of marker genes for airway secretory cells (*Scgb1a1* and *Cyp2f2*) and a pair of neuroendocrine cell markers (*Ascl1* and *Calca*) and mapped them relatively to each other in the bronchiolar epithelium (Figure 6B). In total we analyzed 15 genes encoding soluble secreted proteins (*Scgb1a1*, *Napsa*, *Lgi3*, *Calca*, *Sftpc* and *Lyz2*), cell surface proteins and receptors (*Cd74*, *Cldn18*, *Fgfr2* and *Ager*), a metabolic enzyme *Cyp2f2*, and signaling proteins and transcription factors (*Axin2, Spry2*, and *Etv5*, *Ascl1*) (Figure 6A, B). We first focused on *Sftpc-CreER*^pos^-*Rosa-Ai14*^pos^ AT2 cells and on the analysis of 11 selective AT2 markers covering a broad spectrum of expression levels (Figure 6 A, C). To evaluate the utility of SCRINSHOT in complementing scRNA-Seq data with spatial information we compared the SCRISNSHOT analysis with available scRNA-Seq data from 156 AT2 cells (47). The mean values of SCRINSHOT dots in AT2 cells were proportional to the scRNA-Seq raw count values of AT2 cells. Spearman correlation analysis showed a strong correlation between the results of the two methodologies (ρ=0.9455) for highly-, moderately- and lowly-expressed genes (Figure 6C). As expected, there was no correlation of the SCRINSHOT data from 5615 *Sftpc*^neg^ alveolar cells when we compared them to the AT2 scRNA-Seq dataset (ρ=0.1636). The proportional mean gene expression levels in SCRISHOT and scRNA-Seq argue that SCRINSHOT provides a suitable alternative for rapid, *in situ* evaluation of cell states detected by scRNA-Seq.

**Figure 6.**
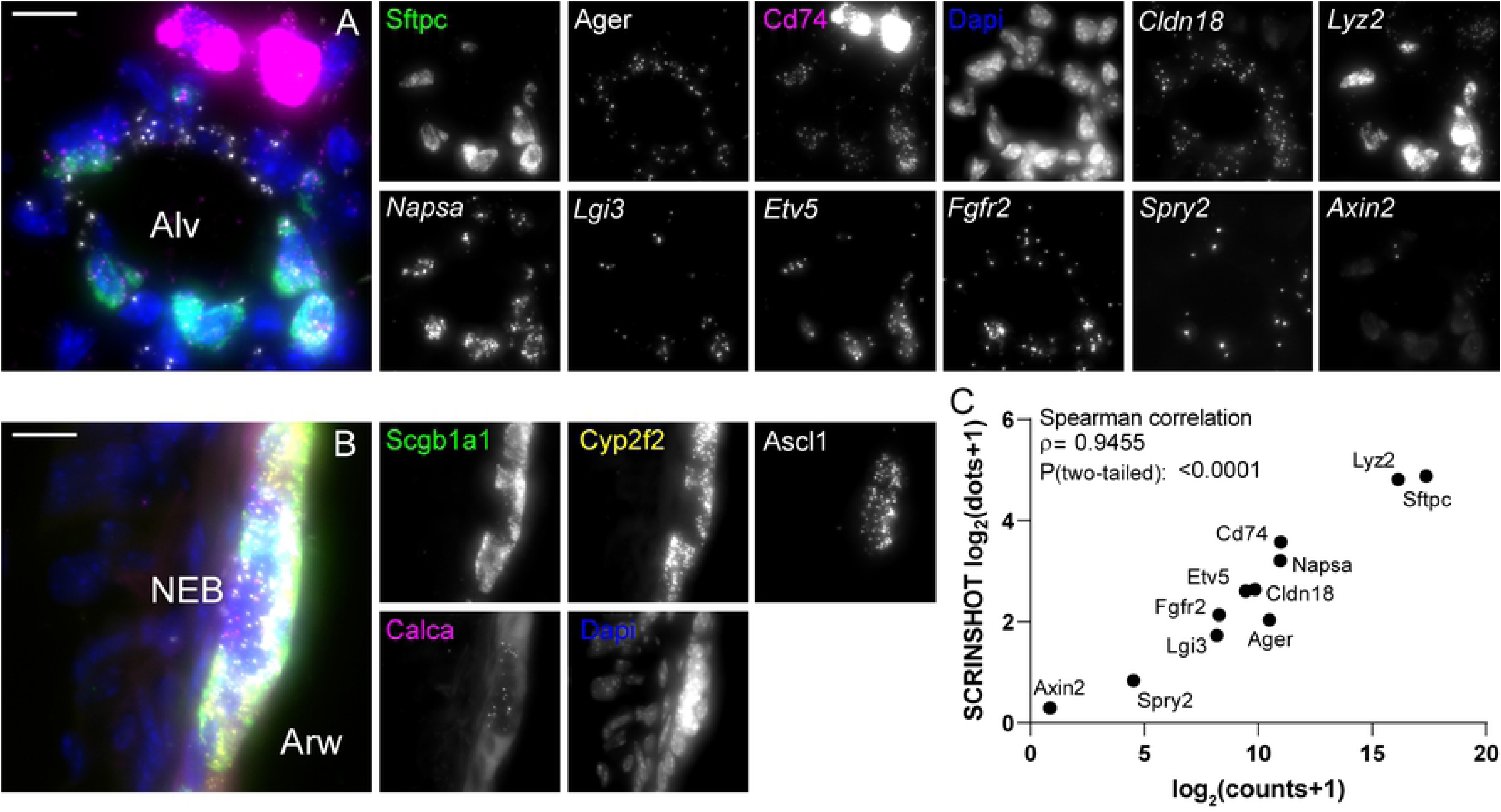
SCRINSHOT multiplexity and correlation with single-cell mRNA sequencing. (A) Representative image of an alveolar region (Alv), showing the SCRINSHOT signal-dots of 11 AT2 selective genes. Merge image includes *Ager* (gray), *Sftpc* (green), *Cd74* (magenta) and DAPI (blue) for nuclear staining. The rest of the genes are indicated with italics and only single-channel images were used. (B) Image of an airway region (arw) with a neuroepithelial body (NEB), containing SCRINSHOT dots for the club-cell markers *Scgb1a1* (green) and *Cyp2f2* (yellow) and neuroendocrine (NE)-cell markers *Calca* (magenta) and *Ascl1* (gray). DAPI (blue) used for nuclear staining. (C) Correlation plot of the indicated genes, between the log_2_(raw counts+1) values of 156 AT2 cells from Liu et al., 2019 and SCRINSHOT signal dots of 1679 AT2 cells. Scale bar: 10μm.

### Spatial mapping of tracheal cell heterogeneity using SCRINSHOT

Recently, two studies addressed the heterogeneity of tracheal epithelium using sc-RNA Seq. These studies identified a new pulmonary cell type, which expresses *Cftr* and therefore considered to play a role in cystic fibrosis pathophysiology (48, 49). They also provided the detailed transcriptomic state of additional cell types, like basal, tuft and secretory cells, including club and two classes of goblet cells.

We tested the ability of SCRINSHOT to detect the above cell types and analyze tracheal epithelial cell heterogeneity with spatial, single-cell resolution. We used a panel of selective markers for club, goblet, basal, tuft and ionocytes, as identified in (48, 49). The 29 analyzed genes included (i) *Scgb1a1, Scgb3a1, Il13ra1, Reg3g, Lgr6* and *Bpifb1* as club-cell markers, (ii) *Foxq1, Gp2, Pax9, Spdef, Tff2, Lipf, Dcpp3* and *Dcpp1* as goblet-cell markers, (iii) *Trp5, Il25, Gng13, Six1, Alox5ap* and *Sox9* as tuft-cell markers, (iv) *Foxi1, Tfcp2l1, Cftr* and *Ascl3* as ionocyte markers, (v) *Ascl1* as a neuroendocrine cell marker (50) and (vi) *Krt5, Pdpn* and *Trp63* as basal cell markers. We also included *Muc5b* as a general secretory marker of the proximal airways (48, 49). For assignment of cell positions we utilized structural landmarks that separate the tracheal airway epithelium in three parts, the proximal, which extends until the end of the submucosal gland, the intermediate part, which spans eight cartilage rings deeper and the distal, which includes the remaining part of trachea epithelium, up to bronchial bifurcation (carina). We also assigned positions to proximal intra-lobar airway epithelial cells, which extend up to the L.L3 branching point (51) and to distal airway epithelial cells located at terminal bronchioles (Figure 7A).

**Figure 7.**
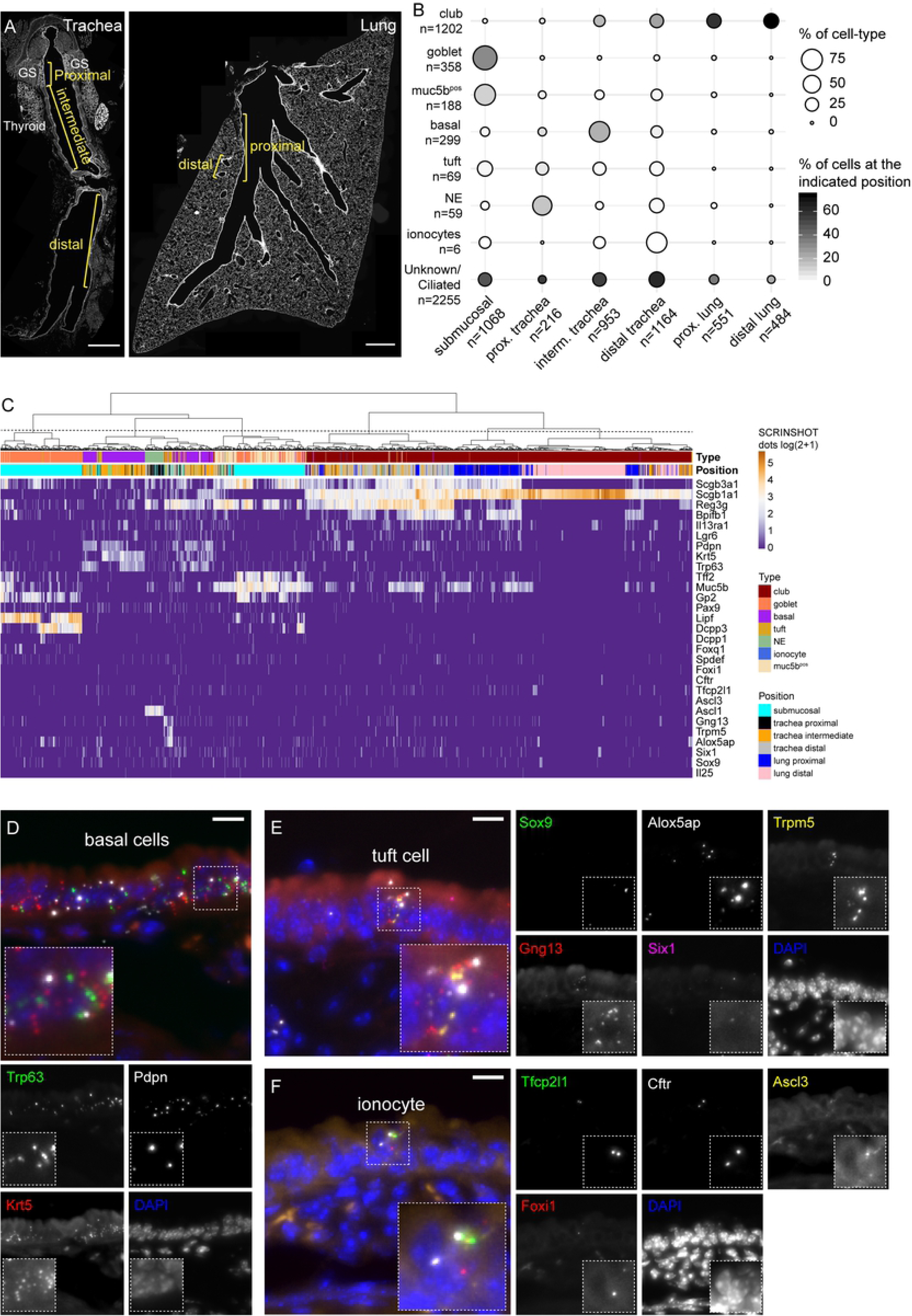
Spatial mapping of tracheal cell types with SCRINSHOT. (A) Overview of analyzed trachea and lung tissue sections using nuclear staining. Analyzed areas are indicated by brackets, corresponding to proximal trachea, submucosal glands (GS), intermediate and distal trachea, followed by proximal and distal lung airways. Trachea image includes also thyroid glands. Scale bar: 500μm. (B) Balloon plot of the annotated cell-types at the analyzed positions. The balloon size indicates the percentage of the cell-type at the indicated position relatively to the total number of cells of the cell-type. The balloon intensity corresponds to the percentage of the specified cell-type, relative of the total number of cells at the indicated position. (C) Hierarchical clustering of the annotated cells. Heatmap shows the log_2_(dots/cell + 1) gene values in the analyzed cells. (D) Characteristic example of basal cells in tracheal epithelium. *Trp63* (green), *Ktr5* (red), *Pdpn* (gray) and DAPI (blue). Scale bar: 10μm. (E) Detection of a tuft cell in tracheal epithelium. *Sox9* (green), *Gng13* (red), *Alox5ap* (gray), *Trpm5* (yellow), *Six1* (magenta) and DAPI (blue). Scale bar: 10μm. (F) An ionocyte in trachea airway epithelium, detected with *Tfcp2l1* (green), *Foxi1* (red), *Cftr* (gray), *Ascl3* (yellow) and Dapi (blue). Scale bar: 10μm.

For cell-type annotation, we initially applied the following threshold criteria for the selected marker genes. We considered club cells, only the *Scgb1a1^pos^* cells, which were negative for the neuroendocrine cell marker *Ascl1* and expressed up to one goblet, tuft, ionocyte and basal-cell markers. *Ascl1^pos^* cells were considered as neuroendocrine cells. Similarly, we annotated the analyzed cells as goblet, basal, tuft cells and ionocytes, if they were positive for at least two characteristic genes of the respective type and only expressed up to one marker of the others. The previously described similarities in gene expression between goblet and club cells in (48), prompted us to consider a cell as goblet if it was positive for at least two of the identified goblet cell markers, regardless of *Scgb1a1* expression. We sampled 1068 cells in submucosal glands and 216 proximal trachea airway epithelial cells. In the intermediate part of the tracheal tube, we quantified 953 cells and 1164 in the distal. In the intra-lobar airway epithelium, we analyzed 551 cells in proximal and 484 in distal airways. SCRINSHOT detected all the previously described trachea cell-types. Club cells comprised 7% of total cells in proximal trachea and their proportion gradually increased to 77% towards the distal intra-lobar airways (Figure 7B). *Trp63^pos^*, *Krt5^pos^* and *Pdpn^pos^* basal cells were primarily detected in the intermediate part of the trachea (21% of the measured cells) and became reduced towards the intra-lobar airways (Figure 7 B, C, D). Tuft cells expressing *Trmp5*, *Gng13 and Alox* were exclusively found in the tracheal epithelium (Figure 7 B, C, E). Ionocytes present a rare cell-type (<1% of trachea epithelium) implicated in the pathogenesis of cystic fibrosis. We detected sparse ionocytes expressing *Cftr* along with the transcription factors *Tfp2l1*, *Ascl3* and *Foxi1* (48, 49) in the tracheal airway epithelium (Figure 7 B, F) and even more rarely in the submucosal glands (Supplementary Figure 6). Their restricted positioning in the tracheal and submucosal gland epithelium highlights the importance of these lung regions in cystic fibrosis caused by *Cftr* mutations in experimental models and in patients (52, 53).

The majority of the *Ascl1^pos^* NE-cells were detected in the proximal-trachea, but as expected we also detected positive cells, scattered along the airways (Figure 7 B, C). Interestingly, 97% of the goblet cells were detected in the submucosal gland but not airway epithelium (Figure 7 B, F). In a hierarchical clustering of all annotated cells, the goblet cells were grouped together with the Muc5b^pos^ cells into 2 clusters (Figure 7C). In agreement with the identification of 65 goblet cells in the sc-RNA Seq data of Montoro et al. (48), *Gp2* is detected in the majority of goblet cells. We also detected *Tff2^pos^ Muc5b^pos^* cells corresponding the described *goblet-1* sub-cluster and cells corresponding to the *Dcpp3^pos^ Lipf^pos^ goblet-2* sub-cluster. Interestingly, SCRINSHOT revealed some additional spatial heterogeneity of *goblet-2* cells in the submucosal glands. We also noticed Gp2^pos^positive cells, expressing high levels of either *Dcpp3* or *Lipf* (Supplementary Figure 5 A) and second, a small subset of the Dccp3^pos^ cells also expressed *Dcpp1* (Supplementary Figure 5 B, arrowhead) (48). The spatial analysis of epithelial cell types in the trachea and lung airways demonstrate the utility of SCRINSHOT in the localization of rare cell types in a complex tissue.

### Spatial mapping of airway and alveolar cells

We used the expression values of 15 genes, in 14167 cells, to generated a spatial map of macrophages, AT1 and AT2 cells in the alveolar compartment and club and neuroendocrine cells in the airways (Supplementary Figure 7). We annotated an airway cell as secretory if it was *Scgb1a1*^pos^ *Cyp2f2*^pos^ *Ascl1*^neg^ and neuroendocrine if it expressed *Ascl1*. In the alveolar compartment of the *Sftpc-CreER*^pos^*-Rosa-Ai14*^pos^ lung, we annotated 1679 *Sftpc*^pos^ cells as AT2. More than 99% of them also contained more than 3 dots of *Lyz2*, another AT2 marker (54), which is also expressed by lung inflammatory cell-types (36, 55). 91.8% of the AT2 cells were also scored positive for *Cd74*. This gene is significantly enriched in AT2 cells but it is also expressed in hematopoietic lineages (56). Additionally, SCRINSHOT analysis distinguished a group of 588 *Lyz2*^pos^ *Cd74*^pos^ *Sftpc*^neg^, which are probably macrophages. We also detected 312 cells positive only for *Lyz2* and 450 expressing only *Cd74*. This suggests that the combinatorial expression of 3 genes defines four cell populations in the alveolar compartment, AT2 cells, *Lyz2*^pos^ *Cd74*^pos^ macrophages and single *Lyz2*^pos^ or single *Cd74*^pos^ immune cells. For identification of AT1 epithelial cells, we used the expression of *Ager* (57). We scored 1129 alveolar *Ager*^pos^ *Sftpc*^neg^ *Scgb1a1*^neg^ cells as AT1. This analysis argues that SCRINSHOT can readily distinguish known epithelial cell types in the lung airways and alveoli. Additional markers for other characterized cell-types, such as endothelial cells and fibroblasts, could be also included in future analyses to facilitate the creation of complete spatial maps of cells types in various organs and help to elucidate gene-expression and cell-type distribution patterns in healthy and diseased tissues.

## CONCLUSION

SCRINSHOT simplifies multiplex, *in situ* detection of RNA in various tissues. The assay is optimized for high sensitivity and specificity. We present a robust analysis protocol with minimal requirements for extensive instrumentation or computational skills, which makes it user-friendly and accessible in many fields of biology. The protocol is suitable for validation of scRNA-seq results in small- or large-scale experiments or for the analysis of gene expression changes upon genetic or chemical manipulations. Importantly, SCRINSHOT is also a low-cost method compared to other *in situ* hybridization methods or antibody detection and the straight forward analysis protocol makes it suitable for routine laboratory use. Although SCRINSHOT multiplexity is reduced compared to “*in situ* sequencing” by ligation (24) or other barcode-based approaches (4, 58), the number of detection fluorophores or hybridization detection cycles can be increased to detect 50 genes per slide. Manual cell segmentation is the most time-consuming part of the analysis and we believe that automatic segmentation will provide a future solution. At present, our attempts with open access programs like Ilastik (59) or Cell Profiler (60) only scored a 50% success rate in segmentation of lung sections presumably because of the elongated and overlapping cell shapes and the large cavities of the lung alveolar compartment. Confocal microscopy and higher magnifications may improve resolution but cost will also increase. The SCRINSHOT analysis of the mouse submucosal gland, trachea and proximal-lung airway epithelium for 29 genes and distal-lung for the expression of 15 genes demonstrates the capacity of the method to detect not only low abundance mRNAs but also rare cell-types, like ionocytes, tuft and neuroendocrine cells, facilitating the generation of cellular maps from complex tissues.

## MATERIALS AND METHODS

### Animals and Histology

All experiments with *wild type* (C57Bl/6J) mice were approved by the Northen Stockholm Animal Ethics Committee (Ethical Permit numbers N254/2014, N91/2016 and N92/2016).

For transgenic mouse experiments, *Sftpc-CreER*^negative^(42);*Rosa26-Ai14*^positive^(44) and *Sftpc-CreER*^negative^;*Rosa26-Ai14*^positive^ individuals were used according to German regulation for animal welfare at the Justus Liebig University of Giessen (Ethical Permit number GI 20/10, Nr. G 21/2017). The recombination and *Sftpc*^pos^ cell labelling of the *Sftpc-CreER*^pos^;*Rosa26-Ai14*^pos^ lung was done by one subcutaneous injection of Taxamoxifen, on postnatal day 1 (P1) and the tissues were collected on P21. An *Sftpc-CreER*^neg^;*Rosa26-Ai14*^pos^ littermate treated as above and used as negative control, in addition to a *wild type* P60 lung.

For lung tissue collection, the mice were anesthetized with a lethal dose of ketamine/xylazine. Lungs were perfused with ice cold PBS 1X pH7.4, through the heart right ventricle to remove red blood cells from the organ. A mixture (2:1 v/v) of (PFA 4% (Merck, 104005) in PBS 1X pH7.4):OCT (Leica Surgipath, FSC22) was injected into the lung, from the trachea using an insulin syringe with 20-24G plastic catheter B Braun 4251130-01), until the tip of the accessory lobe got inflated and the trachea was tied with surgical silk (Vömel, 14739). The lungs were removed and placed in PFA 4% in PBS 1X pH7.4 for 4 hours for P21 lungs and 8 hours for adult, at 4°C in the dark, with gentle rotation or shaking. The other analyzed organs were simply placed in PFA 4% in PBS 1X pH7.4 for 8 hours.

The tissues were transferred to a new tube with a mixture (2:1 v/v) of (30% sucrose in PBS 1X pH7.4):OCT at 4°C for 12-16 hours, with gentle rotation or shaking. Thereafter, tissues were embedded in OCT (Leica Surgipath, FSC22), using specific molds (Leica Surgipath, 3803025) and frozen in a slurry of isopentane and dry ice. Tissue-OCT blocks were kept at −80°C until sectioning. 10 μm thick sections were cut with a cryostat (Leica CM3050S) and placed on poly-lysine slides (Thermo, J2800AMNZ), kept at room temperature for 3 hours with silica gel (Merck, 101969) and then stored at −80°C for further use.

### Embryonic human lungs

Use of human fetal material from elective routine abortions was approved by the Swedish National Board of Health and Welfare and the analysis using this material were approved by the Swedish Ethical Review Authority (2018/769-31). After the clinical staff acquired informed written consent by the patient, the tissue retrieved was transferred to the research prenatal material lung sample was retrieved from a fetus at 8.5 w post-conception. The lung tissue was fixed for 8 hours at 4°C and processed as above.

### Probe Design

A detailed description of the Padlock probe design is provided in the Additional File 1. Briefly, the PrimerQuest online tool (Integrated DNA Technologies: IDT) was used to select sequences of Taqman probes (40-45 nucleotides) for the targeted mRNA. These sequences were then interrogated against targeted-organism genome and transcriptome, with Blastn tool (NLM) to guarantee their specificity. The Padlock Design Assistant.xlsm file was used to split the sequences in two and integrate them into the padlock backbone. Padlock probes were ordered from IDT as Ultramer DNA oligos and their sequences are provided in Additional File 2.

The 40-45 Taqman probe sequences were also used to prepare the fluorophore-labelled oligos. Using the IDT OligoAnalyzer tool the length of the sequences was adjusted to Tm 56°C. To remove the fluorescent oligos after each detection cycle, we exchanged “T” nucleotides with “U” and treated with Uracil-DNA Glycosylase (Thermo, EN0362). Detection oligos were labelled at their 3’-end with fluorophores and manufactured by Eurofins Genomics. The sequences of fluorophore-labelled oligos are provided in Additional File 3.

### Pretreatments of the slides

Slides were transferred from −80°C to 45°C to reduce moisture. A post fixation step with 4% PFA in PBS 1x pH7.4 was done, followed by washes with PBS-Tween20 0.05%. Permeabilization of tissues was done with 0.1M HCl for 3 minutes, followed by two washes with PBS-Tween20 0.05% and dehydration with a series of ethanol. SecureSeal^TM^ hybridization chambers (Grace Bio-Labs) were mounted on the slides and sections were preconditioned for 30 minutes at room temperature (R/T) with hybridization-reaction mixture of 1X Ampligase Buffer (Lucigen, A1905B), 0.05M KCl (Sigma-Aldrich, 60142), 20% deionized Formamide (Sigma-Aldrich, F9037), 0.2μg/ul BSA (New England Biolabs, B9000S), 1U/μl Ribolock (Thermo, EO0384) and 0.2μg/μl tRNA (Ambion, AM7119). To block unspecific binding of DNA we included 0.1μM Oligo-dT30 VN.

### Padlock probe hybridization, ligation and RCA

Hybridization of the Padlock probes was done in the above solution, omitting Oligo-dT30 VN and adding 0.05 μM of each Padlock probe. We used three padlock probes for every targeted RNA-species. For the highly abundant *Scgb1a1* mRNA, we used 0.01 μM of one padlock probe to minimize molecular and optical saturation. Hybridization included a denaturation step at 55°C for 15 min and an annealing step at 45°C for 2 hours. Not hybridized padlock probes were removed by washes with 10% Formamide in 2X SSC (Sigma-Aldrich, S6639). To minimize the effect of the previously documented SplintR ligase nucleotide preferences (26, 33), padlock probe ligation was performed overnight O/N at 25°C using the SplintR ligase (NEB, M0375) at a final concentration of 0.5 Units/μl, T4 RNA ligase buffer (NEB, B0216) and 10μM ATP, according to manufacturer recommendations (see New England Biolabs webpage).

Rolling Cycle Amplification (RCA) was done O/N at 30°C using 0.5 Units/μl Φ29 polymerase (Lucigen, 30221-2). The reaction mixture contained also 1X Φ29 buffer, 5% Glycerol, 0.25 mM dNTPs (Thermo, R0193), 0.2 μg/μl BSA and 0.1 μM RCA primers (RCA Primer1: TAAATAGACGCAGTCAGT*A*A and RCA Primer2: CGCAAGATATACG*T*C). The “*” indicate Thiophosphate-modified bounds to inhibit the 3-5 exonuclease activity of Φ29 polymerase (23). A fixation step with 4% PFA for 15 minutes was done to ensure stabilization of the RCA-products on the tissue. Sections were thoroughly washed with PBS-Tween20 0.05% before next step.

### Hybridization of detection-oligos

The visualization of the RCA products was done with hybridization of the 3’-fluorophore labeled detection oligos. The reaction mixture contained 2X SSC, 20% deionized Formamide, 0.2 μg/μl BSA, 0.5 ng/μl DAPI (Biolegend, 422801), 0.4 μM FITC-labeled and 0.2 μM Cy3- and Cy5-labeled detection probes for 1 hour at R/T. Washes were performed with 10% Formamide in 2X SSC, followed by 6X SSC. Tissues sections were dehydrated with a series of ethanol, chambers were removed and the slides were covered with SlowFade™ Gold Antifade Mountant (Thermo, S36936) and a coverslip.

After image acquisition, the coverslips were removed by placing the slides in 70% ethanol in 45°C. Then, sections were dehydrated using a series of ethanol to mount the hybridization chambers. After tissue rehydration with PBS-Tween20 0.05%, the detection-oligos were digested with Uracil-DNA Glycosylase for 1hour at 37°C. The reaction mixture contained 1X UNG buffer, 0.2 μg/μl BSA and 0.02 Units/μl Uracil-DNA Glycosylase (Thermo, EN0362). Destabilized oligos were stripped off by thorough washes with 65% deionized Formamide at 30°C. Multiple rounds of hybridization and imaging, as described above, were performed until all genes were imaged.

### Image acquisition

Images were captured with a Zeiss Axio Observer Z.2 fluorescent microscope with a Colibri led light source, equipped with a Zeiss AxioCam 506 Mono digital camera and an automated stage, set to detect the same regions after every hybridization cycle.

### Image analysis

The nuclear staining was used to align the images of the same areas between the hybridizations and multi-channel *.czi files, containing the images of all genes, were created using Zen2.5 (Carl Zeiss Microscopy GmbH). The images were analyzed as 16bit *.tiff files, without compression or scaling. Images were tiled using a custom script in Matlab (The MathWorks, Inc.). Manual nuclear segmentation was done with Fiji ROI Manager (61). The nuclear ROIs were expanded for 2 μm with a custom Cell-Profiler script and considered as cells. The signal-dots were counted in these cell-ROIs using Cell-Profiler 3.15 (60), Fiji (62, 63) and R-RStudio (58, 64–67) custom scripts (https://github.com/AlexSount/SCRINSHOT).

### Thresholding

A cell-ROI size criterion was applied to remove the outliers with very small or big surface. In particular, only cells included between two standard deviations of the mean size of the analyzed cells were further processed. In SCRINSHOT we considered a cell-ROI positive for an analyzed gene, we used a threshold strategy. First, we determined the maximum numbers of signal-dots per cell-ROI for all analyzed genes. A cell-ROI was considered as positive, if it contained more than 10% of the maximum number of signal-dots for the specific gene. The higher threshold was set to 3, which was applied for highly abundant genes with maxima over 31 signal dots.

### Curation of the data

In general, the 2 μm nuclear expansion provides an underestimation of real signal dots and provides satisfactory results for airway cells (e.g. Figure 3A’ merge-images with cell-ROI outlines) but the cellular segmentation of the alveolar region is more challenging mainly because of the irregular cell shapes and their overlap. This gave false positive cell-ROIs due to dots from adjacent true-positive cells being erroneously assigned to their neighbors. To reduce the noise, signal-dots of highly abundant RNA-species were used to visually inspect and remove the problematic cell-ROIs from further analysis.

### Clustering

Annotated cells of submucosal gland, trachea and lung airway epithelium were clustered using hclust package in R (68). Log_2_(dots/cell + 1) values were used to calculate Euclidean distances and clustering was done using ward.D2 method. Balloon plots were created by ggpubr package in R (69) and heatmaps with pheatmap package (70). Bootstrapping analysis was done by the clusterboot package (71). Clusters were considered as “stable” if bootstrap values were >0.5.

### Analysis of *Sftpc-CreER*^pos^;*Rosa26-Ai14*^pos^ cells

For the identification of the RFP^pos^ alveolar cell-ROIs in the *Sftpc-CreER*^pos^;*Rosa26-Ai14*^pos^ lung, all analyzed alveolar cells from the RFP^neg^ *Sftpc-CreER*^neg^;*Rosa26-Ai14*^pos^ lung were used to determine the maximum endogenous fluorescence (in Raw Integrated Density values) and set it as a threshold. The RFP^pos^ and RFP^neg^ cell-ROIs were curated and the *Sftpc* signal-dots were measured in them.

For the correlation of RFP endogenous fluorescence with SCRINSHOT *RFP* signal, the endogenous fluorescence (in Raw Integrated Density values) of all segmented cell-ROIs from the analyzed tissue sections was correlated with the detected RFP signal-dots in the same cell-ROIs, using simple linear regression analysis in Graphpad Prism with default settings.

### Comparison of SplintR and cDNA-based detection of RNA species

To compare the performance of padlock-probe hybridization (i) directly on RNA (SCRINSHOT) and (ii) after cDNA synthesis, we used sequential 10μm-thick sections from adult mouse lungs, fixed for 8 hours (see above). cDNA-based approaches in earlier publications have only used fresh frozen tissues and pepsin or proteinase K tissue treatments to increase RNA accessibility (3, 24). The padlock probes were designed to recognize exactly the same sequence of the analyzed genes. For the highly (*Scgb1a1, Sftpc*) and intermediate (*Actb*) expressed genes, we used only one padlock probe and for *Pecam1*, three probes because of its lower expression levels.

For cDNA synthesis, the RNA species of the tissue sections were transformed to cDNA by reverse transcription (RT) using random decamers. The RNA strands were degraded with RNaseH to let padlock probes hybridize to the corresponding cDNA sequences. Padlock probe ligation was done with Ampligase DNA ligase. All other steps were done according to the provided SCRINSHOT protocol.

For cDNA synthesis we used 20 U/µl of the SuperScript™ II Reverse Transcriptase (Thermo, 18064014), 1X SuperScript™ II RT buffer (Thermo), 0.5 mM dNTPs (Thermo, R0193), 10mers-random primer (Thermo), 0.2 µg/µl BSA (NEB, B9000S), and 1 U/µl RiboLock RNase Inhibitor (Thermo, EO0384), at 42°C, O/N. The slides were post-fixed with 4% (w/v) PFA in PBS 1X pH7.4, at RT for 30 min, following by 6 washes with PBS-Tween 0.05%.

### Evaluation of SCRINSHOT specificity using mutated padlock probes

The experiment was done as described above, using 0.01 μM of *Scgb1a1* padlock probes and 0.05 μM of the *Actb*. The “mismatch” probe had the same sequence as the normal *Scgb1a1* padlock probe with a C>G substitution at the 5’-ligation site and the “3’-scrambled” probe had the same 5’-arm but the 3’-arm was scrambled. *Actb* padlock probe was used as internal control to calculate the *Scgb1a1/Actb* ratios. The *Scgb1a1/Actb* ratios of cell-ROIs with zero *Actb* signal-dots were considered as zeros. The detection of all *Scgb1a1* RCA-products was done using a detection oligo, which recognizes the padlock-backbone of *Scgb1a1* but not of *Actb*. *Actb* RCA-product was detected by a detection oligo, which recognizes its gene-specific sequence (Additional File 3). The *Scgb1a1/Actb* fluorescence ratios were calculated using the Raw Integrated Densities of the two genes in each cell-ROI.

### Evaluation of SCRINSHOT specificity using an antisense competitor

To compete the binding of Scgb1a1 padlock probe on its target transcript, an antisense competitor (A), which recognizes the Scgb1a1 mRNA between nucleotides 316-378 and masks the padlock probe hybridization site, was used in the same hybridization mixture with the *Scgb1a1* padlock probe (P) at different ratios: (i) P:A=1:0, (ii) P:A=1:1, (iii) P:A=1:5, keeping the *Scgb1a1* padlock probe concentration to 0.05μM.

### Correlation with SCRINSHOT with published single cell RNA sequencing datasets

The GSE118891 dataset was used to retrieve gene expression values (raw counts) of all AT2, according to the cell annotation of provided metadata file (47). The genes of interest were selected and their Mean values were calculated and log_2_(dots+1) transformed. Similarly, the SCRINSHOT signal-dots per cell-ROI were log_2_(dots+1) transformed. Pearson correlation analysis was done using GraphPad Prism.

### Statistical analysis

All statistical analyses were done with Graphpad Prism, using nonparametric tests, since the SCRINSHOT data do not follow canonical distributions. Multiple comparisons were done using ANOVA Kruskal-Wallis multiple comparison test, without multiple comparison correction (“*”: P ≤ 0.05, “**”: P ≤ 0.01, “***”: P ≤ 0.001, “****”: P ≤ 0.0001). For pairwise comparisons the statistical analysis was done using Mann-Whitney nonparametric t-test (“*”: P ≤ 0.05, “**”: P ≤ 0.01, “***”: P ≤ 0.001, “****”: P ≤ 0.0001). Spearman correlation was used to examine the correlation between SCRINSHOT and scRNA-Seq data.

### Availability of data and materials

The datasets and analysis files of the current study have been deposited at Zenodo repository (DOI: 10.5281/zenodo.3634561). All scripts are available at https://github.com/AlexSount/SCRINSHOT.

## Author contributions

CS and AS conceived the idea and designed the experiments with help by MN and WS. ABF and AF provided mouse tissues and ES the human fetal material. AS and HPN performed all hybridizations. AS, AL and XQ wrote the analysis scripts. AS, AL and ABF analyzed the datasets. All authors helped interpreting results. CS, AS and HPN wrote the manuscript and AS the detailed protocol (Additional file 1) with input from all other authors. All authors read and approved the final manuscript. CS provided all funding support.

## Acknowledgments

We would like to thank the Associate Professor Qi Dai for the thorough reading and suggestions of the detailed SCRINSHOT protocol (Additional File 1).

---

**Supplementary Figure 1.**
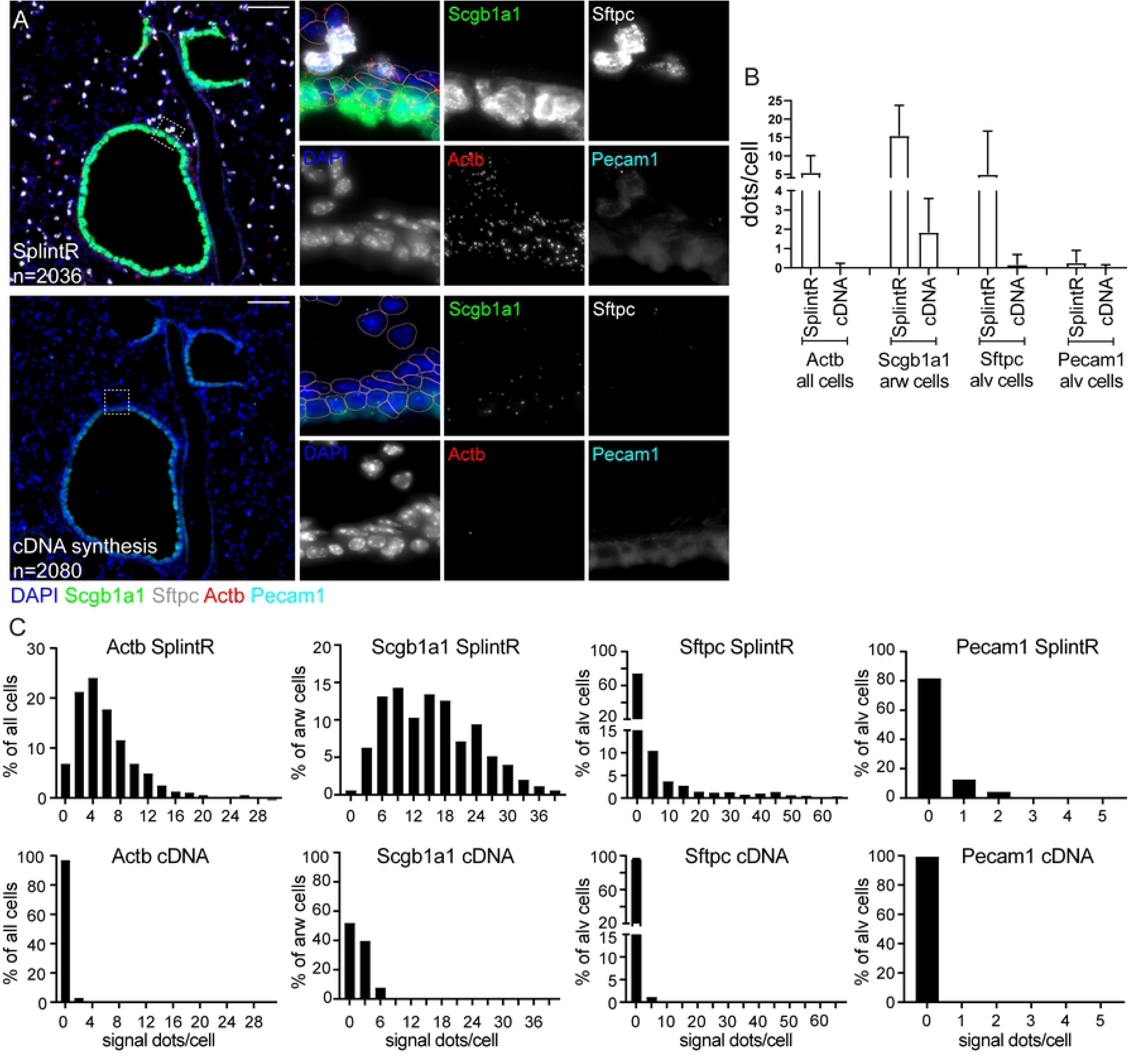
Comparison of SplintR-based (SCRINSHOT) and the cDNA-based *in situ* hybridization assays for high, intermediate and low abundant genes in sequential PFA fixed lung sections. (A) Images of SplintR-based (SCRINSHOT) and cDNA-based *in situ* hybridization assays, in sequential lung sections. DAPI: blue, *Scgb1a1*: green, *Sftpc*: gray, *Actb*: red and *Pecam1*: cyan. Pink outlines show the 2 μm expanded airway nuclear ROIs, which are considered as cells. The square brackets indicate the magnified areas on the right. The “n” correspond to the number of counted cells in large images. Scale bar: 100μm. (B) Bar-plots of the analyzed gene signals, in the indicated tissue compartments, for SCRINSHOT and cDNA-based approaches. The differences between the two conditions are significant (P < 0.0001) for all analyzed genes. (C) Histograms of the analyzed genes. The Y-axes indicate the percentage of the cell-ROIs and the X-axes, the binned signal dots in each cell. In SplintR-condition, 350 cells localized in airways (arw) and 1624 in alveolar (alv) compartment. In cDNA-condition, there are 295 airway and 1706 alveolar cells. Analysis was done using raw images, with the same acquisition conditions and thresholds. Only for visualization purposes, signal intensity of *Scgb1a1* and *Sftpc* in cDNA-condition was set 5-times higher than SplintR.

**Supplementary Figure 2.**
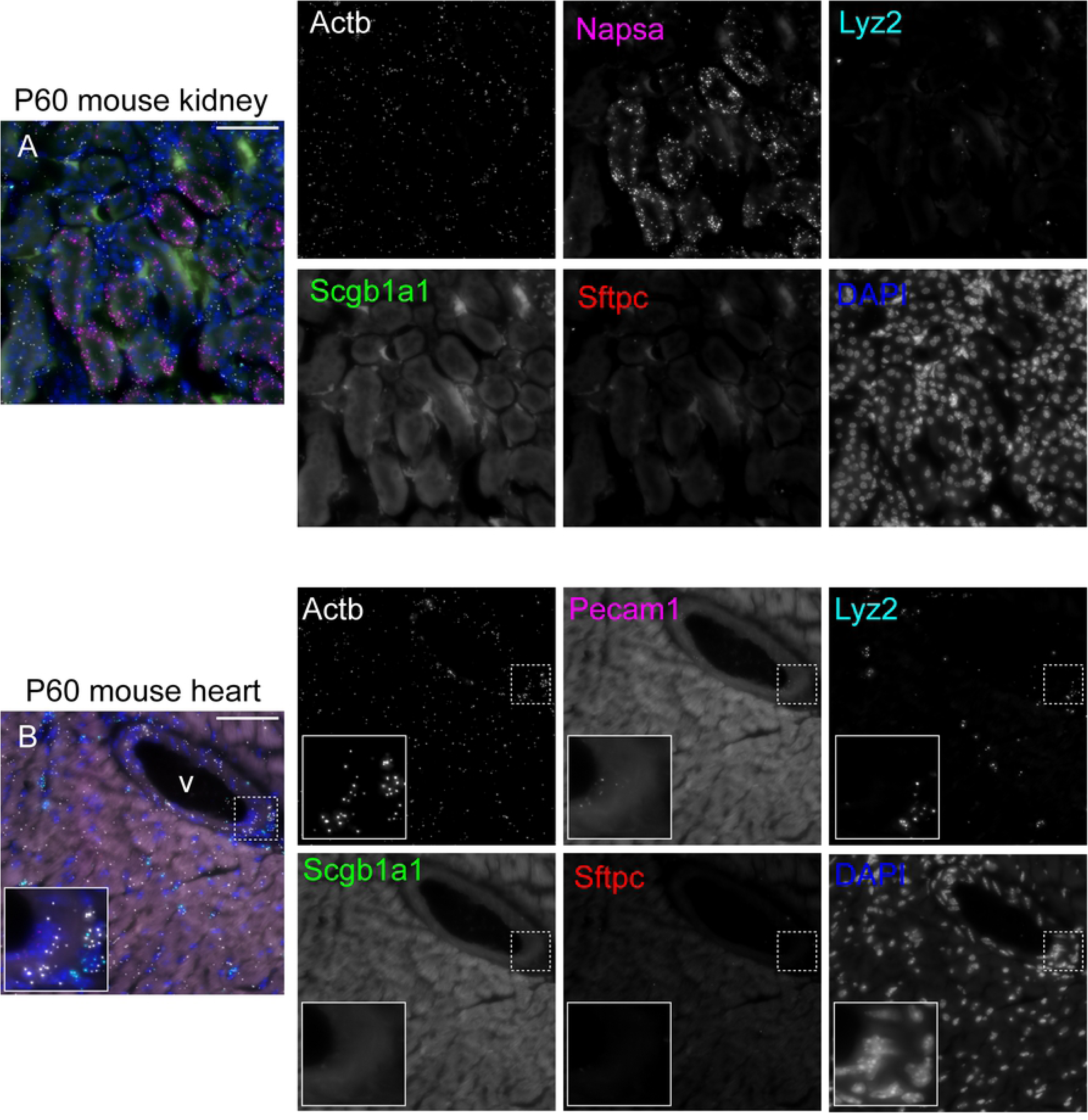
SCRINSHOT application on mouse kidney and heart. (A) Application of SCRINSHOT in adult mouse kidney sections shows signal for *Actb* (gray), *Napsa* (magenta) and *Lyz2* (cyan) but not *Scgb1a1* (green) and *Sftpc* (red). (B) Representative image from an adult mouse heart section, containing a vessel (v). *Pecam1* (magenta) is detected at the vessel walls, *Actb* (gray) is uniformly expressed and *Lyz2* (cyan) labels a few cells (presumably macrophages) but not *Scgb1a1* (green) and *Sftpc* (red) are not detected. DAPI: blue, scale bar: 50μm.

**Supplementary Figure 3.**
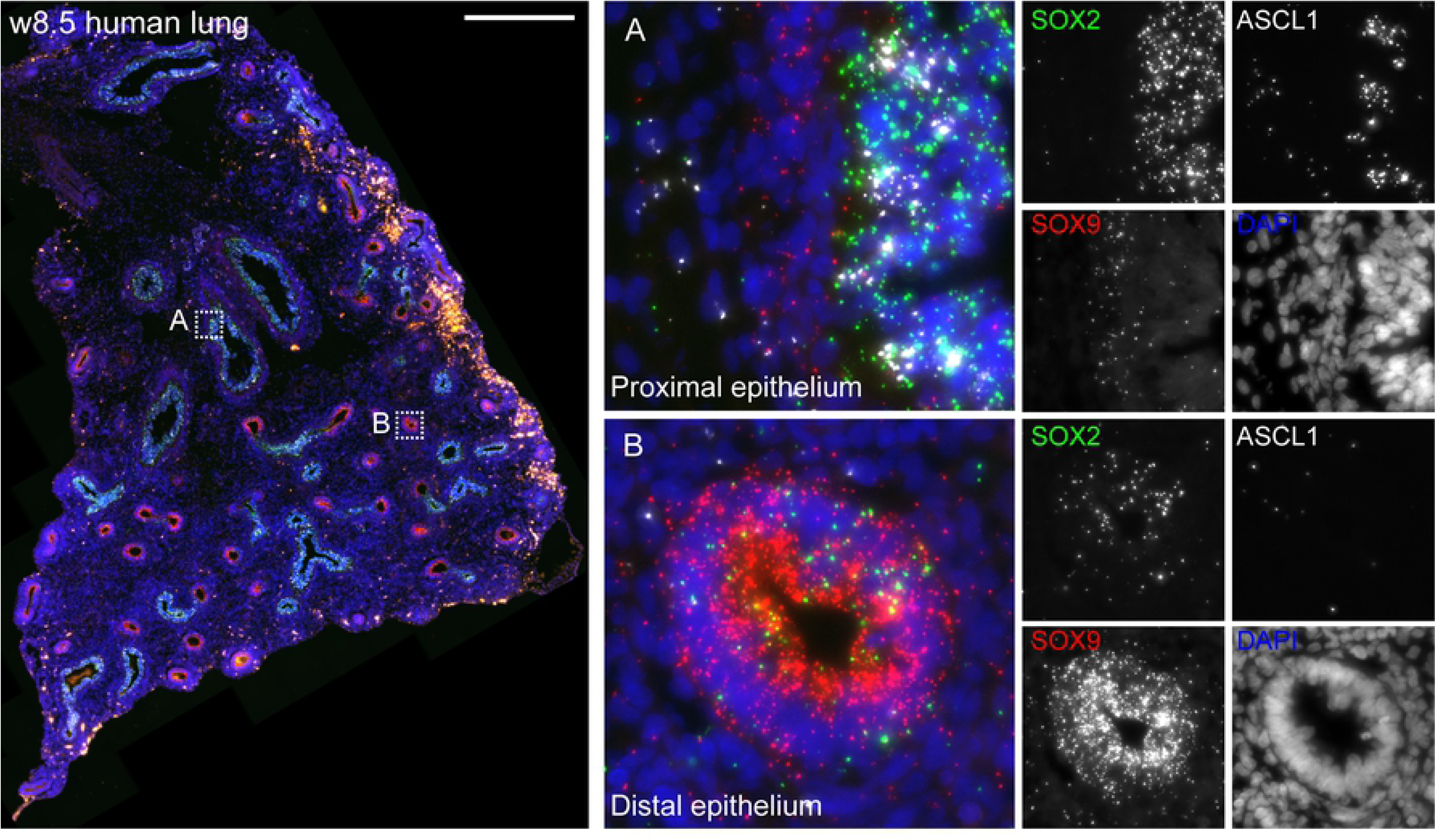
SCRINSHOT application on fetal human lung. On the left, overview of a w8.5 whole left lung tissue section, showing SCRINSHOT signal for *SOX2* (green), *SOX9* (red) and *ASCL1* (gray). The square brackets correspond to the images on the right. DAPI (blue) was used as nuclear staining. Scale bar: 500μm. (A) Representative image of proximal epithelium, which is highly positive of *SOX2* and *ASCL1* but not *SOX9*. (B) Representative image of highly *SOX9* positive distal epithelium.

**Supplementary Figure 4.**
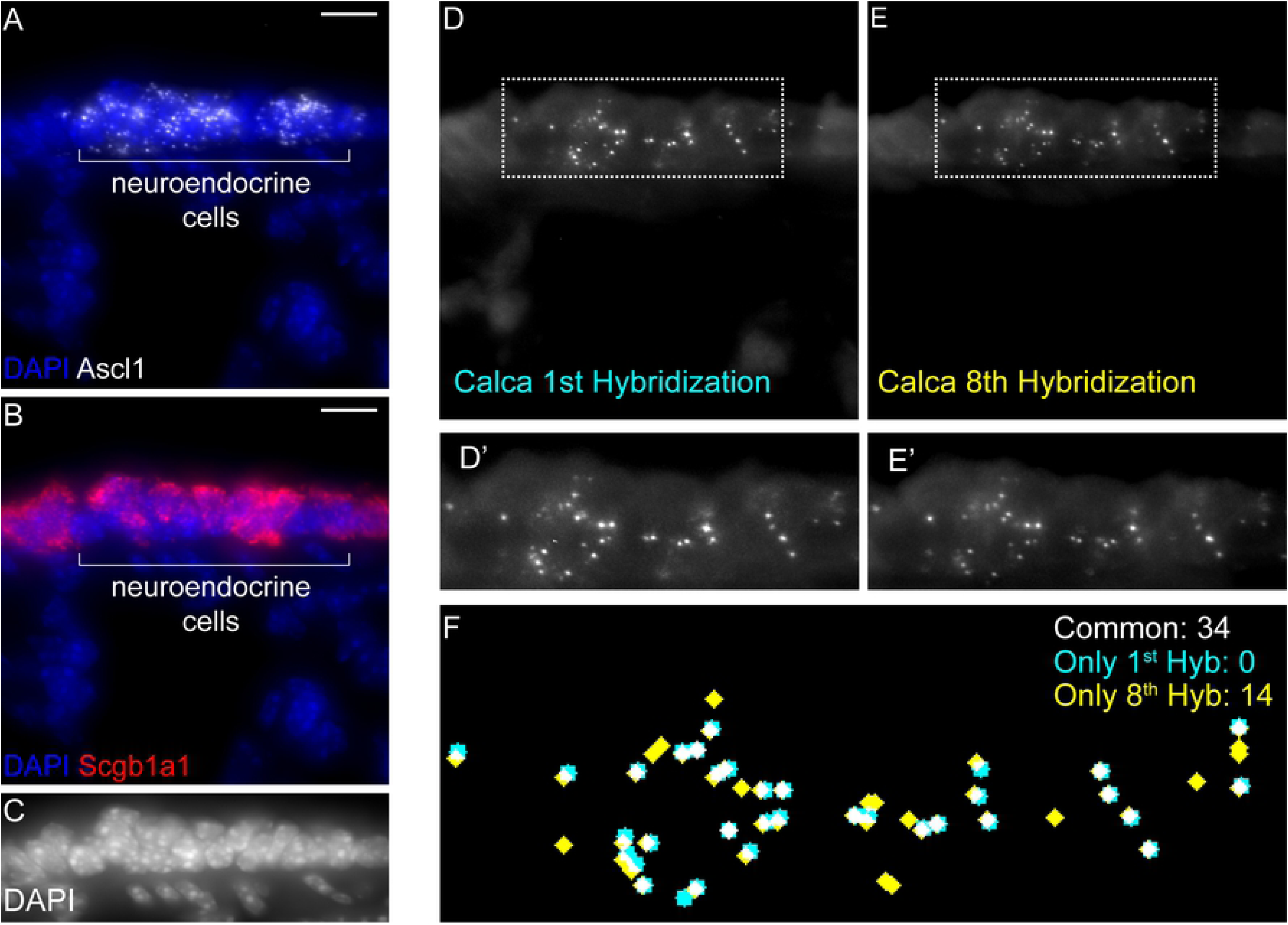
Comparison of detected *Calca* RCA-products in first and eighth hybridizations. (A, B) Images on the left show the *Ascl1*^pos^ (gray) neuroendocrine cells of an airway neuroepithelial body, in relation to *Scgb1a1*^pos^ (red) club cells. DAPI: blue, scale bar: 10μm. (C) Note that neuroepithelial bodies are tightly packed cellular structures, as indicated by DAPI nuclear staining. The images on the right show the *Calca* RCA-products, detected in the first (D) and the eighth (E) detection cycles. (D’-F’) Magnified areas of the indicated positions (brackets) of images D-F, respectively. (F) Overlay of *Calca* identified signal-dots in first (cyan) and eighth (yellow) detection cycles, using the same threshold in CellProfiler.

**Supplementary Figure 5.**
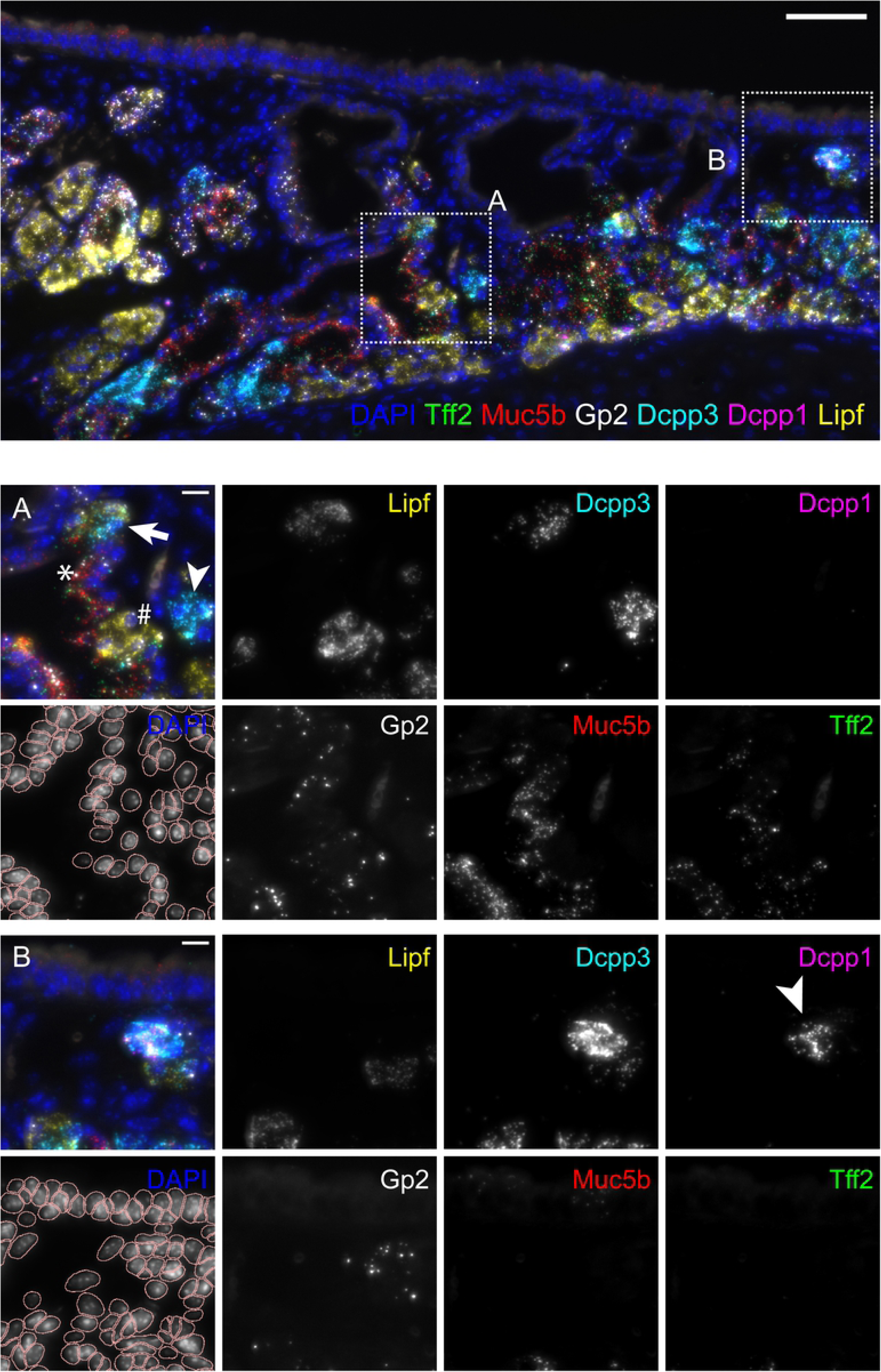
SCRINSHOT mapping of submucosal glands reveals spatial heterogeneity in goblet cell population. Overview of the analyzed submucosal gland for the expression of 6 goblet cell markers shows their expression in submucosal gland but not airway epithelium. *Muc5b* is detected along the airway epithelium of the proximal trachea, indicating that it is a general proximal epithelial cell marker. *Tff2*: green, *Muc5b*: red, *Gp2*: gray, *Dcpp3*: cyan, *Dcpp1*: magenta, *Lipf*: yellow, DAPI: blue and cell-ROIs: pink. Scale bar: 500μm. (A) Insert showing the previously described *Muc5b^pos^ Tff2^pos^* goblet subtype (arrow) and the *Lipf^pos^ Dcpp3^pos^* (asterisk). *Lipf^pos^ (hash)* and *Dcpp3^pos^* (arrowhead) are detected in the same region, being positive for the general goblet cell marker *Gp2*. (B) Insert showing regionally restricted expression of *Dcpp1* in a subset of *Dcpp3^pos^* cells. Insert scale bar: 10μm.

**Supplementary Figure 6.**
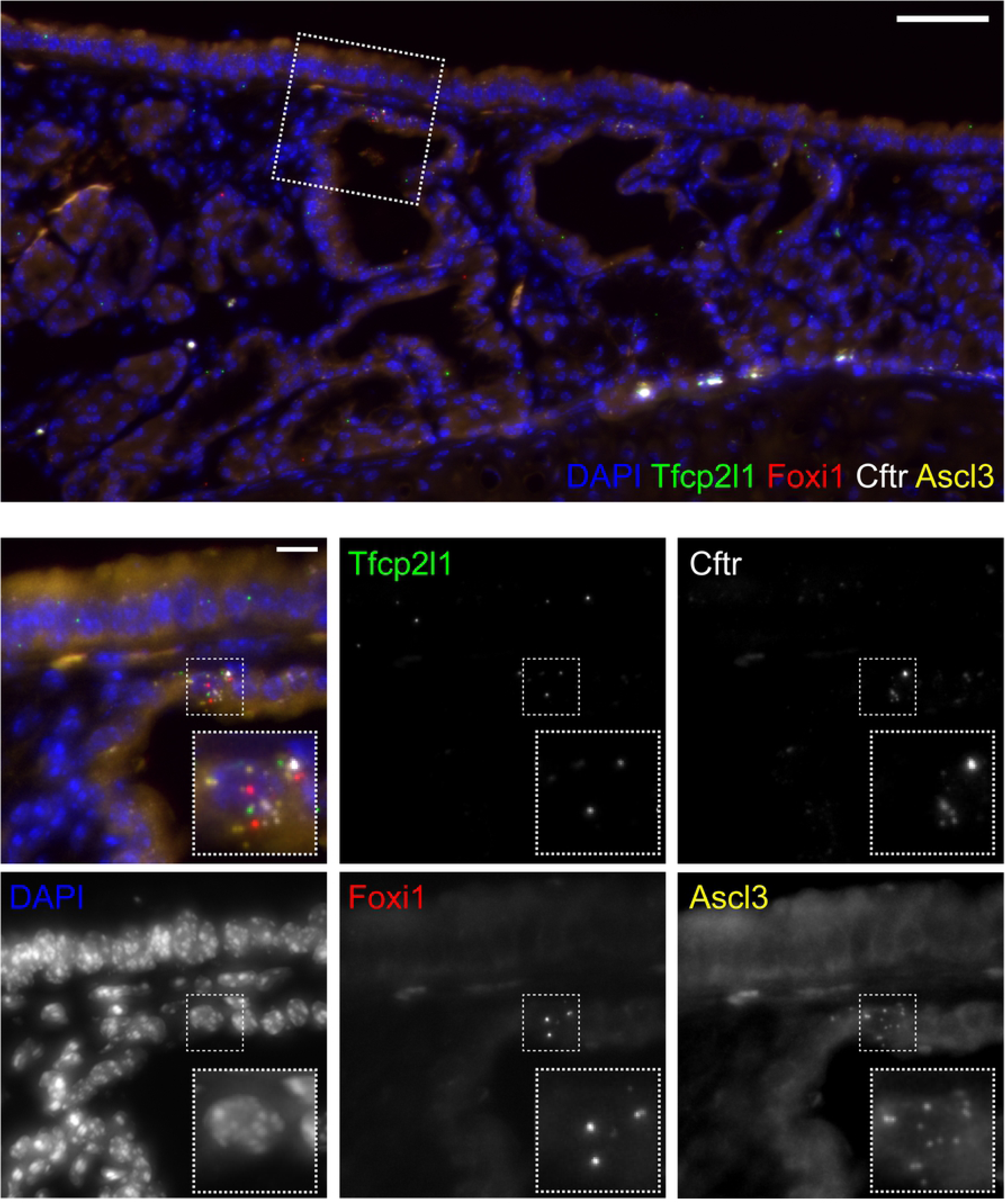
Ionocyte in the submucosal gland. (Top) overview of the analyzed submucosal gland, showing SCRINSHOT signal for ionocyte markers, *Tfcp2l1* (green), *Cftr* (gray), *Foxi1* (red), *Ascl3* (yellow) and DAPI (blue). Scale bar: 500μm. (Bottom) Magnified area of the indicated square in overview image, showing a detected ionocyte in submucosal gland. Scale bar: 10μm.

**Supplementary Figure 7.**
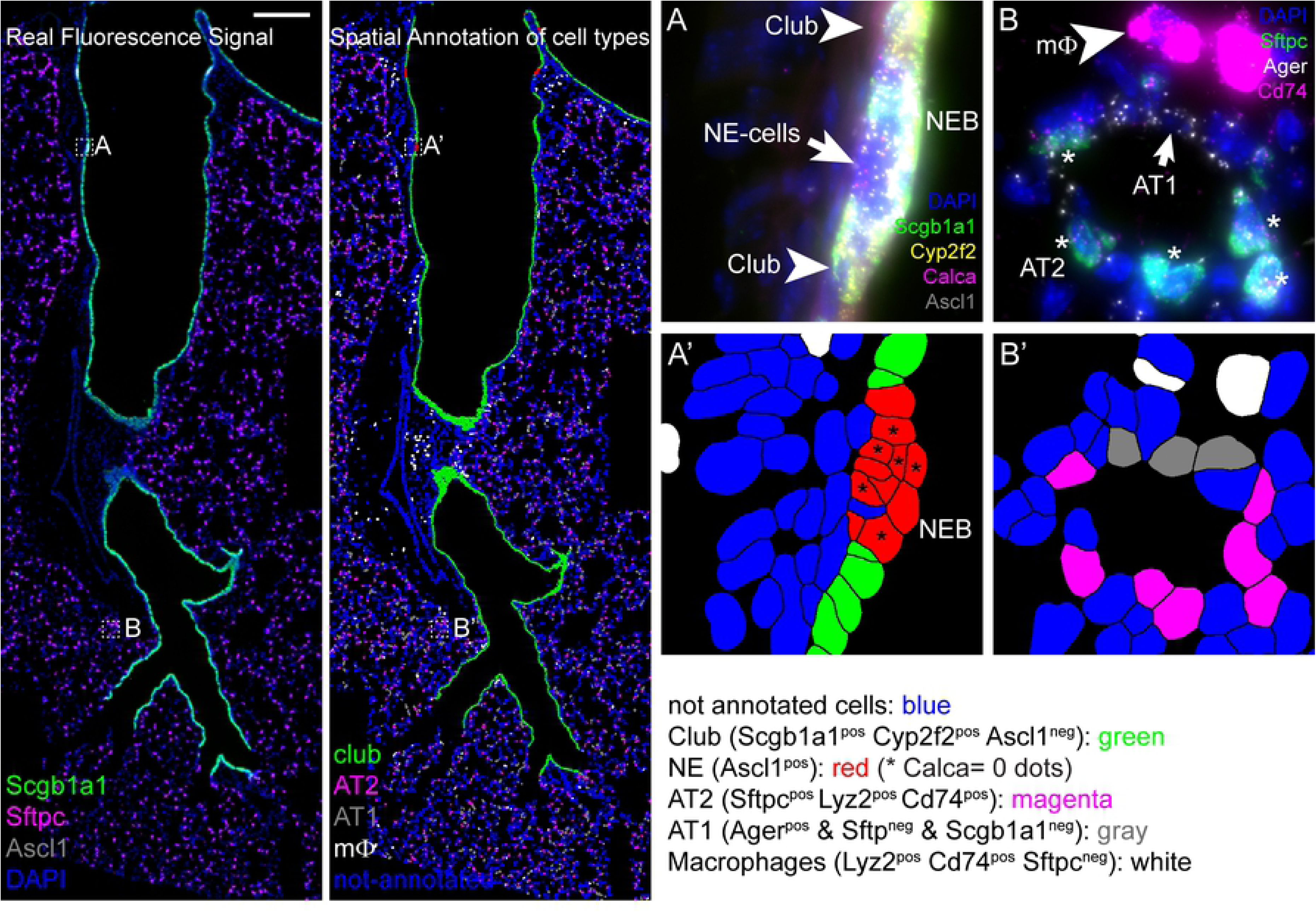
SCRINSHOT generates cell-type digital annotation maps of large tissue areas. (i) Overview image of SCRINSHOT fluorescence signal-dots for *Scgb1a1* (green), *Sftpc* (magenta) and *Ascl1* (gray) of a large area from P21 *Sftpc-CreER*^pos^;*Rosa-Ai14*^pos^ lung section, after Tamoxifen induction on P1 (RFP was not shown at that image). The image contains 14167 manual segmented nuclei, which were expanded for 2 μm and considered as cells. (ii) Spatial map of annotated cell types according to the indicated criteria. (A) Same airway area as in Figure 6B, showing club (*Scgb1a1*: green and *Cyp2f2*: yellow) and NE-cell (*Calca*: magenta and *Ascl1*: gray) markers, in an airway position with a neuro-epithelial body (NEB). (A’) Cell-type digital annotation of the area corresponding to “A”. The “*” indicate *Ascl1^pos^ Calca^neg^* cells Club cells: green and NE-cells: red. (B) Same alveolar area as in Figure 6A showing *Ager^pos^* (gray) *Sftpc*^neg^ (green) *Cd74*^neg^ (magenta) AT1 cells (arrow), *Ager^low^ Sftpc*^pos^ *Cd74^low^* AT2 cells (asterisks) and *Ager^neg^ Sftpc*^neg^ *Lyz2^pos^ Cd74^high^* macrophages (mΦ, arrowhead). (B’) Cell-type digital annotation of the area corresponding to “B”. AT1 cells: gray, AT2 cells: magenta and macrophages: white. All not annotated cells are depicted with blue. Scale bar: 200μm.

